# Migration of endocrine and metabolism disrupting chemicals from plastic food packaging

**DOI:** 10.1101/2024.04.05.588263

**Authors:** Sarah Stevens, Zdenka Bartosova, Johannes Völker, Martin Wagner

## Abstract

Plastics constitute a vast array of substances, with over 16 000 known plastic chemicals, including intentionally and non-intentionally added substances. Toxicity and thousands of chemicals are extractable from plastics; however, the extent to which toxicity and chemicals migrate from everyday plastic products remains poorly understood. This study aims to characterize the endocrine and metabolism disrupting activity, as well as chemical composition of migrates from plastic food contact articles (FCAs) from four countries as significant sources of human exposure. Additionally, strategies for prioritization of chemicals were explored. Fourteen plastic FCAs covering seven polymer types with high global market shares were migrated into water and a water-ethanol mixture as food simulants according to European regulations. The migrates were analyzed using reporter gene assays for nuclear receptors relevant to human health and non-target chemical analysis to characterize the chemical composition of the migrates. All FCA migrates interfered with at least two nuclear receptors, predominantly targeting pregnane X receptor. Moreover, peroxisome proliferator receptor gamma was broadly activated by the migrates, though mostly with lower potencies, while estrogenic and antiandrogenic activities were more selectively induced by specific FCAs. Fewer chemicals and less toxicity migrated into water compared to the water-ethanol mixture. The latter exhibited similar toxicity and number of chemicals as methanol extracts of the same FCAs. Novel strategies were employed to address the chemical complexity of FCAs and narrow down the list of potential active chemicals. By comparing the composition of multiple leachates of one sample and using stepwise partial least squares regressions, we successfully reduced the chemical complexity, pinpointed potential endocrine disruptors such as triphenyl phosphate and prioritized chemicals for further identification efforts. This study demonstrates the migration of endocrine and metabolism disrupting chemicals from plastic FCAs into food simulants, rendering a migration of these compounds into food and beverages probable.

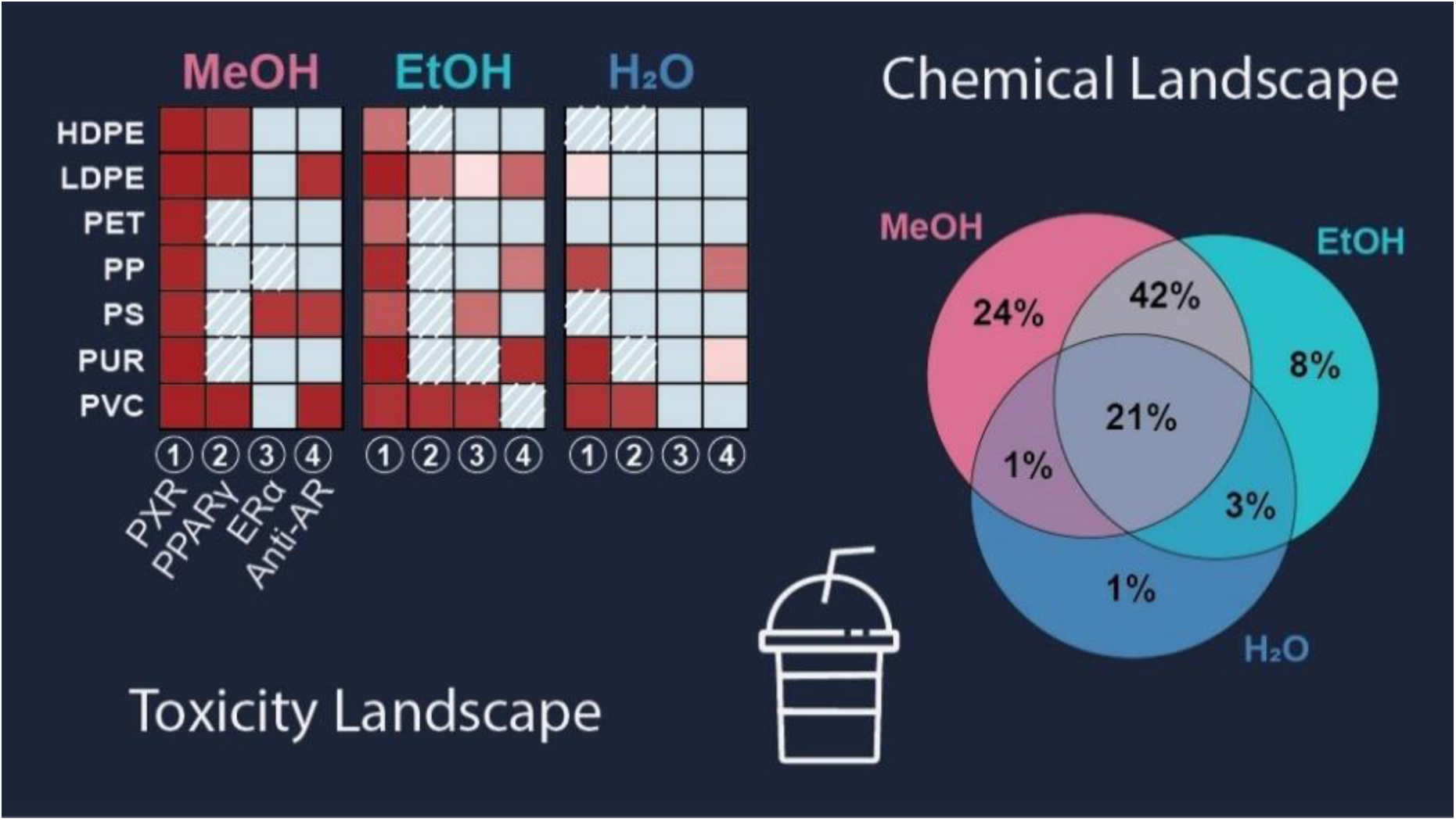

## 1. INTRODUCTION

The annual health costs attributed to a few plastic chemicals are totaling an estimated $249 billion in the United States alone.^1^ These estimates are based on a few well studied plastic chemicals, such as polybrominated diphenylethers, phthalates, bisphenols, and polyfluoroalkyl substances and perfluoroalkyl substances. However, plastics constitute a much vaster array of chemicals, with a recent study identifying more than 16 000 known plastic chemicals, encompassing both chemicals used in production and non-intentionally added substances (NIAS).^2–4^ While one quarter of these chemicals is of concern, almost two thirds remain poorly studied, lacking essential hazard data^4^ and additional plastic chemicals remain unidentified, with a significant portion probably being NIAS.^5,6^

Plastic chemicals which are largely non-covalently bound to the polymer, can be released from plastic products, and contaminate human and natural environments. Humans can be exposed to these chemicals through various routes, including direct dermal or oral contact, inhalation of volatile substances or contaminated dust. Exposure via ingestion, that is, an exposure to chemicals migrating from food contact articles (FCA), represents a prevalent route.^7^ The migration of plastic chemicals into food is influenced by factors such as temperature, contact duration, chemical properties, and the properties of the food or beverage.^8^ Regulatory testing of FCAs relies on food simulants that mimic properties like hydrophilicity, lipophilicity and acidity of foods, with different duration and temperature regimes applied to cover various use scenarios.^9^

Once humans are exposed to plastic chemicals, they can pose potential health risks by interfering with hormone systems or metabolic processes.^10,11^ Several plastic chemicals are known endocrine and metabolism disrupting chemicals (EDCs and MDCs) that can act via nuclear receptors.^10,12^ However, these represent only a fraction of the chemicals detected in plastic products.^13^ Leachate studies are essential to comprehensively assess the toxicity of the chemical mixtures released from plastic products, integrating the effects of unknown compounds.^14^ In our recent study,^15^ we extracted chemicals from plastic FCAs, revealing widespread in vitro endocrine and metabolism disruptive activity and a large chemical complexity. However, the question remains whether these toxic compounds transfer to foods and thus, are available to human exposure.

A challenge encountered with leachate toxicity studies is the identification of causative compounds, stemming from the large chemical complexity of plastic leachates. In this context, employing data reduction techniques like partial least squares (PLS) regression can serve as a valuable tool to filter for potentially active compounds.^16^ Several methods are available for the selection of variables important in contributing to receptor activity, among them stepwise selection methods based on variable importance of projection (VIP).^17^ This approach has been used previously for identification of chemicals features covarying with toxicity.^15,18,19^

In our previous study^15^, we used methanol as solvent to extract the chemicals from plastic FCAs, representing a worst-case scenario. This study extends our previous approach, evaluating the chemicals and toxicity that are likely to transfer to foods under more realistic conditions. Here we compared the migration of plastic chemicals into water (best-case scenario) and a water-ethanol mixture (1:1, v:v), simulating fatty foods (food simulant D1) in accordance with EU regulations (European Regulation 10/2011/EU).^9^ To assess the metabolism disrupting and endocrine disrupting activity of the plastic migrates, we employed reporter gene assays covering four receptors important for human health: pregnane X receptor (PXR), peroxisome proliferator receptor gamma (PPARγ), estrogen receptor alpha (ERα), and androgen receptor (AR).

Additionally, non-target high resolution mass spectrometry was used for chemical screening and to tentatively identify migrating chemicals. By comparing the toxicity profiles and the different sets of chemicals released from FCAs with various solvents, and using PLS regressions, we further aim to prioritize toxic chemicals for additional investigations. Hereby, we demonstrated that plastic FCAs are a relevant source of exposure to EDCs and MDCs. Further, we tentatively identified known hazardous compounds and prioritized chemical features.

## 2. MATERIALS AND METHODS

### 2.1 Samples

Fourteen plastic FCAs made of the seven polymer types with the highest global market share were purchased from domestic retailers from four countries (Germany, Norway, South Korea and UK, Table 1). The samples comprise a subset of the FCAs used in our previous study, selected to include two samples per polymer type not containing food, except for sample PP 2. The samples were purchased between winter 2020 and spring 2021 and transported to the laboratory in polyethylene (PE) bags (VWR). One FCA, the coffee cup (PP 2), contained foods, which was removed by washing with tap water at the local sites. The polymer type of the FCAs was identified through Fourier-transformed infrared spectroscopy and differential scanning calorimetry for high- and low-density polyethylene (HDPE, LDPE).^15^ If the packaging included information about the polymer type, we used that information to label the products.

**Table 1.** Plastic food contact articles analyzed in this study. Note that the sample names corresponding to the ones used in our previous study^15^ can be found in Table S1.

| polymer | sample name | product | country |
| --- | --- | --- | --- |
| HDPE | HDPE 1 | food container | Germany |
|  | HDPE 2 | freezer bag | South Korea |
| LDPE | LDPE 1 | zip lock freezer bag | Germany |
|  | LDPE 2 | zip lock freezer bag | UK |
| PET | PET 1 | oven bag | Germany |
|  | PET 2 | food container | Norway |
| PP | PP 1 | bowl | South Korea |
|  | PP 2 | coffee cup<sup>a</sup> | South Korea |
| PS | PS 1 | cup | South Korea |
|  | PS 2 | styrofoam tray | Norway |
| PUR | PUR 1 | hydration bladder | Germany |
|  | PUR 2 | hydration bladder | Norway |
| PVC | PVC 1 | drinking tube | Germany |
|  | PVC 2 | cling film | UK |

<sup>a</sup> This item had contact with food prior to analysis. High- and low-density polyethylene (HDPE, LDPE), polyethylene terephthalate (PET), polypropylene (PP), polystyrene (PS), polyurethane (PUR), and polyvinyl chloride (PVC).

### 2.2 Migration

We used ultrapure water (18.2 MΩ cm^-1^, PURLAB flex, ELGA, water migrate) as best case scenario and food simulant D1, a water-ethanol mixture in a 1:1 (v:v) ratio (ethanol ≥ 99.9%, Supelco, water-ethanol migrate) for oil in water emulsions as food simulants for the 10-d migration experiment according to European Regulation 10/2011/EU. To prevent sample contamination, all consumables used in the process (except plastic pipette tips for volumes < 1 mL) were made of glass or stainless steel, rinsed with ultrapure water (UPW) and acetone, and heated at 200 °C for at least 2 h. We used 27 g of each FCA per food simulant, which was cut into smaller pieces (0.5−0.8 × 2 cm, thickness ≤0.4 cm), placed in 250 mL glass bottles (Schott) and added 180 mL food simulant (0.15 g plastic mL^-1^). The bottles were closed with aluminum foil to impede evaporation and randomly placed for 10 d in a heating chamber (Termarks) at 40 °C. Every second day, the samples were gently agitated to ensure that all plastic pieces were in contact with the food simulant. A water saturated environment was maintained by placing trays with UPW in the chamber which were refilled daily. Parallel to the FCAs migration, three procedural blanks (PB) for each solvent not containing plastics were treated identically throughout the whole process.

### 2.3 Solid Phase Extraction

We used TELOS C18(EC)/ENV SPE cartridges (700 mg, 6 mL, 06478-09, Cole-Parmer) to extract the samples after the 10-d migration. In preparation for the SPE, the water-ethanol migrates were diluted to a final of 10% ethanol by adding 640 mL UPW to 160 mL of the migrates to reduce loss of chemicals due to a too high ethanol content. The water migrates were treated identically to ensure similar conditions. The SPE cartridges were conditioned with 2 mL n-heptane (purity ≥99.5%, Supelco), followed by 2 mL acetone (purity >99.9%, Sigma-Aldrich), 6 mL methanol (99.8%, Sigma-Aldrich), and 8 mL UPW by gravity. The cartridges were then loaded with the acidified migrates (pH 2.5, adjusted with 3.5 M sulfuric acid) under a constant vacuum flow of approximately 2–5 mL min^-1^. Subsequently, the cartridges were dried under a nitrogen stream and stored at -20°C. Within 1–2 days, samples were eluted consecutively with 5 mL acetone and 5 mL methanol by gravity, with vacuum applied at the end of the process. The eluted samples were further concentrated to 1 mL under a gentle stream of nitrogen (40°C, Reacti-vap III, Thermo Scientific) and transferred to HPLC vials. Evaporation continued until there was space in the vials, at which point 200 μL of dimethyl sulfoxide (DMSO) was added. The remaining acetone and methanol were then further evaporated until the volume did not decrease further, resulting in 800-fold concentrated samples.

In addition to the migrates, we used methanol extracts of the same samples from our previous study.^15^ Here, we extracted 13 g of each FCA with 90 mL methanol (99.8%, Sigma-Aldrich) for 1 h by sonication (Ultrasonic Cleaner USC-TH, VWR) at room temperature. Subsequently, 60 mL of extracts were evaporated under a gentle stream of nitrogen and transferred to 600 mL DMSO, as described for the migrates. This resulted in 100-fold concentrated extracts.

### 2.4 SPE recovery

To assess the recovery of chemical features in the SPE, the water and water-ethanol migrates of three selected FCAs (HDPE 2, LDPE 1 and PVC 1) and PB 1 were analyzed directly without any further treatment (before SPE), and after SPE, following transfer to DMSO and re-dilution to the migration concentration of 0.15 g mL^-1^. To that aim, we took a 0.1 mL sample directly after the 10-d incubation period, transferred it to HPLC vials and stored it at -80°C until further analysis. Additionally, to assess the impact of ethanol on the performance of the SPE, we prepared two water migrates, combined them after the 10-day migration, and introduced 10% ethanol to half of the water migrates prior to the SPE.

### 2.5 Chemical Analysis

For the chemical analysis of the plastic migrates and extracts, the samples dissolved in DMSO were diluted 100-fold and 800-fold, respectively, with a mixture of UPW and methanol (1:1, v:v), resulting in a plastic concentration of 0.15 g mL^-1^, consistent with the concentration during migration and extraction. Chemical analysis was performed with an ultra-high performance liquid chromatography system (Acquity I-Class UPLC, Waters) coupled to a high-definition hybrid quadrupole/time-of-flight mass spectrometer Synapt G2-S (Waters) as described previously^15^ with minor modifications. The separation was performed on an Acquity UPLC BEH C18 column (150 × 2.1 mm ID, 1.7 μm, Waters) in a 20 min gradient with water and methanol as mobile phases, both containing 0.1% formic acid (Table S2). The injection volume was 1 μL. The mass spectrometer was equipped with an electron spray ionization source and operated in positive and negative mode (Table S3). Data was collected across the mass range of 99−1200 Da using data-independent acquisition (resolution 20 000). See the Supplementary Information for further details (S1.1). For the SPE recovery analysis with the three selected FCAs, a different gradient was applied for separation (details in S1.1). Access to mass spectral data for all samples is available at https://doi.org/10.18710/QMMER5. Features with abundance less than 10 times the highest abundance across all PBs were excluded from the analysis. The abundance of features was further adjusted by subtracting the maximum abundance of the respective features detected in the PBs. All features with an abundance >100 (approximately 25th percentile) were used for further analysis.

### 2.6 Compound Identification and Toxicity Data

For a tentative identification (identification level 3, according to Schymanski et al.^20^), we used the Metascope algorithm in Progenesis QI (Nonlinear Dynamics, version 3.0) to compare the experimental spectra with empirical spectra from MassBank (14 788 unique compounds, release version 2021.03), and spectra predicted *in silico*. For the *in silico* predicted spectra, we constructed a database containing 12 297 plastic chemicals of a previous version of the PlastChem report^4^, proceeding as described in the Supplementary Information (S1.2). For the identification, the spectra of each feature in the samples were compared to those in the database, with a precursor ion tolerance of 5 ppm and a fragment ion tolerance of 10 ppm. Tentative identifications were filtered for hits with a score of ≥40, selecting the highest score in cases of multiple identifications. Toxicity data of the tentatively identified compounds detected across all three solvents or otherwise prioritized, were retrieved from the latest ToxCast and Tox21 databases (INVITRODB_V4_1_SUMMARY, US EPA).^21^ We extracted activity concentrations 50 (AC_50_) from 39 assays corresponding to the receptors analyzed here (Table S5). Further, the PlastChem report^4^ was consulted for additional hazard and use information of prioritized compounds.

### 2.7 Reporter Gene Assays

To analyze the receptor activity of the FCA migrates, we used CALUX reporter gene assays (BioDetection Systems B.V., Amsterdam) for human pregnane X receptor (PXR), peroxisome proliferator-activated receptor γ (PPARγ), estrogen receptor α (ERα), and androgen receptor (AR). The assays were performed and analyzed as described in Stevens et al.^15^ In short, each plate included negative controls (assay medium), vehicle controls (assay medium with 0.2% DMSO), and a concentration series of the respective reference compounds (PXR: nicardipine, PPARγ: rosiglitazone, ERα: 17 beta-estradiol, AR: flutamide, Table S6 and Figure S1). The AR assay was operated in antagonistic mode with 0.5 μM dihydrotestosterone (CAS 521-18-6, Sigma-Aldrich) as a background agonist (corresponds to EC_80_ in agonistic mode, see Figure S2). Plastic migrates were 500-fold diluted in assay medium and analyzed in five 1:2 serial concentrations, with the highest concentration containing chemicals that migrated from 12 mg plastic well^-1^. Cytotoxic samples, defined as a 20% reduction in cell count compared to the pooled controls, were further diluted at a 1:2 ratio until reaching non-cytotoxic concentrations. After 23 h of exposure, high-content imaging was used to assess cytotoxicity and normalize the reporter gene response. Receptor activity was analyzed by measuring luminescence (Cytation 5) of the lysed cells for 1 s following injection of 30 μL illuminate mix containing D-luciferin as a substrate. Migrates were analyzed in at least three independent experiments, each with four technical replicates.

For the *in vitro* activity of the methanolic FCAs extracts, we used the data generated in our previous study.^15^ The extracts were analyzed at 8–16-fold lower concentrations, with 1.5 mg plastic well^−1^ (PPARγ, ERα, and AR) or 0.75 mg plastic well^−1^ (PXR) as the highest analyzed concentration. The tested concentrations of the migrates and extracts overlapped in 1 or 2 concentrations for non-cytotoxic samples.

### 2.7 Data Analysis Reporter Gene Assays

We used GraphPad Prism (v10, Graph Pad Software, San Diego, CA) and Microsoft Excel for Windows (v2021−2306) for analyzing the bioassay data as described previously.^15^ In short, we used a four-parameter logistic regression to fit the dose-response relationship of the samples and reference compounds. Receptor activity was expressed as luminescence normalized to the number of cells well^−1^ and further normalized to the dose-response relationship of the reference compound analyzed on the same plate. The activity threshold (limit of detection, LOD) was set to three times the standard deviation of the pooled negative and solvent controls.

### 2.8 Partial Least-Squares Regression

To select chemical features covarying with receptor activity, we conducted stepwise PLS regressions using the R package mdatools.^22^ By this stepwise approach the chemical complexity of the models can be reduced, and models with the best performance can be selected.^23^ We followed a similar approach as in our previous study^15^ and described in Hug et al.^18^ A separate model was established for each receptor. To be able to include extracts and migrates which were tested at different concentrations, we included only methanol extracts either exhibiting or lacking an EC_20/50_ in both, extracts and migrates (Table S7). For migrates with activity > LOD < 20% (50% for antiandrogenic activity), EC values were set to 12 mg well^-1^, corresponding to highest tested concentration of non-toxic samples. The reciprocal of the EC values was use in the models so that larger values were indicative of more potent activity. Inactive samples were set to zero. A stepwise variable selection method based on the variables’ importance of projection (VIP) was employed, excluding features with a VIP < 0.8. The best performing model was selected based on the cross-validated model parameter. Model validation was performed with 500 iterations of a random set with the same number of features as in the optimized models.

## 3. RESULTS AND DISCUSSION

### 3.1 Chemical analysis

#### Recovery of chemical features in the SPE process

The impact of sample preparation on the number of features is rarely investigated in non-target studies of plastic leachate. Hence, we compared the recovery of features in three samples before and after SPE, for results of the procedural blanks see Supplementary Information (S 2.1). We investigated the influence of 10% EtOH on the SPE given that this was the final concentration used to extract water-ethanol migrates. While most features (83–86%) were detected in water migrates with and without 10% EtOH, 11–23% of features were exclusively detected in the migrates with 10% EtOH (Table S9). It is unlikely that these are impurities originating from the ethanol or the SPE cartridge as all features present in the PBs were removed from the analysis. However, reactions between the plastic chemicals and EtOH, such as ethanolysis, can lead to the generation of artefacts.^24^ To further assess the recovery of features after sample preparation, we compared directly injected samples with those subjected to SPE and transferred to DMSO. We found large differences in the recovery between the six samples, with a loss of up to 74% (median 31%) of the chemical features during SPE (Table S10, Figure S3, Figure S4). Moreover, 0.5–76% (median 11%) were added in the extraction process. The largest fraction of features was lost from the HDPE freezer bag (HDPE 2), of which we recovered only 26–28% of the features migrating into water and water-ethanol. Such a loss might occur, among others, due to the selectivity of the cartridge and/or the volatilization of compounds in the evaporation step.^25,26^ This can lead to an underestimation of toxicity.^27^ However, a preconcentration of the plastic migrates is required to compensate for the dilution in the bioassays. Most additional features were detected in the water migrates of LDPE 1 and PVC 1, with 49% and 76%, respectively. The introduction of chemicals can potentially lead to an overestimation of toxicity. However, by analyzing PB along with the samples this is controlled for and the enrichment of environmental samples via SPE is a standard procedure.^27,28^

#### Chemical landscape of the FCAs

Across all FCA extracts and leachates, we detected a total of 17 581 chemical features in positive and 5974 in negative ionization mode. Given the substantially lower number of chemicals detected in negative ionization (Figure 1C), we focus on the results obtained in positive ionization mode here (further details in Table S11 and S17). When considering all samples, we found that 11 262 out of 15 430 extractable chemical features also migrated into food simulants (Figure 1A). This means that 73% of the chemicals present in plastic FCAs leach into food simulants. In addition, 2153 chemical features migrated into water or water-ethanol but were not extracted with methanol. This shows that additional chemicals can leach into food simulants that are not readily extractable.

**Figure 1.**
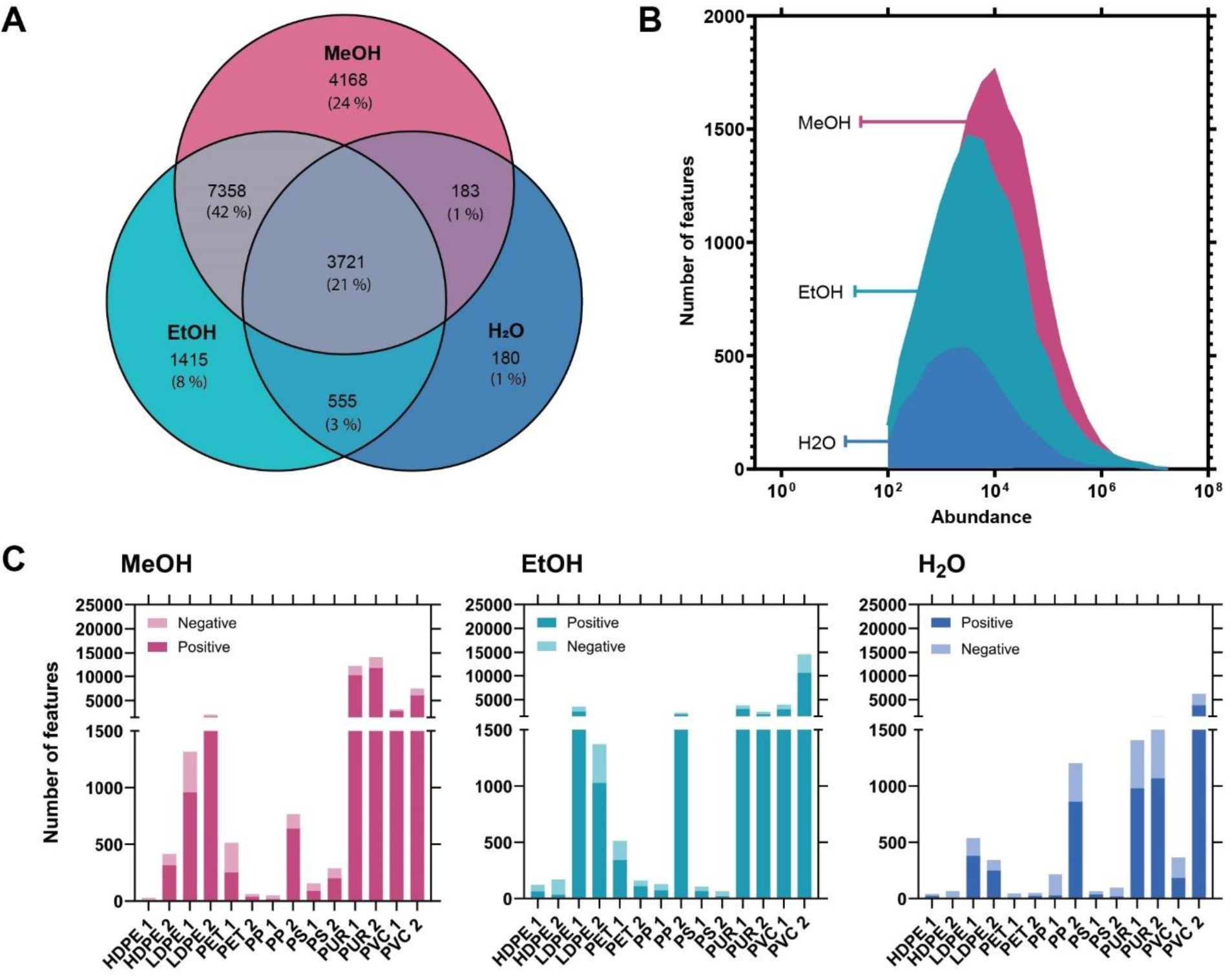
Overlap of chemical features leaching into methanol (MeOH), water-ethanol mixtures (50% EtOH) or water (H_2_O, A), abundance of chemical features per solvent (B), and comparison of number of detected features between the different solvents and ionization modes (C). Note: All features with an abundance <100 were removed from the analyses.

We found large differences in the number of chemical features between the individual FCAs, ranging from 6–11 751 for methanol extracts, 20–10 631 for water-ethanol migrates and 8–3812 for water migrates (Table S11). While the number and abundance of features extracted with methanol was somewhat higher, it is rather similar to water-ethanol migrates (Figure 1A and B). Surprisingly, we detected more features in the water-ethanol migrates than in methanol extracts in eight out of fourteen samples (Figure S5, Table S11). While the largest fraction of all features (42%, 7358 features) was detected in methanol and water-ethanol, 3721 features (21%) were detected in all three solvents rendering human exposure to these chemical features most likely (Figure 1A). As expected, fewer chemicals migrated into water. However, a minor fraction of chemical features (1%) was present only in the water migrates, comprising up to 253 features unique to a single sample (cling film PVC 2, Figure S5). These results demonstrate that most chemicals migrate from plastic food packaging into food simulants and, hence, become available for human exposure. Furthermore, the notion that methanol extracts more chemicals than a food simulant and is, thus, unrealistic, cannot be confirmed for several water-ethanol migrates.

In general, the number of features detected in methanol extracts correlates well with those in both migrates (Spearman r = 0.73–0.79), indicating that products containing more extractable chemicals also leach more chemicals into food simulants (Figure S6). Despite this correlation, a surprisingly large heterogeneity was detected in the chemical composition of the three samples derived from a single FCA (Figure S5). As an example, we detected a distinct set of features in each sample of the HDPE freezer bag (HDPE 2) and the PET food container (PET 2) (Figure S5B and F). Moreover, both PS products leached unique chemical features into water, different to those migrating into water-ethanol or extracted with methanol (Figure S5I and J). This suggests that the PS products contain a small set of hydrophilic chemicals found only in the water migrates, while most chemicals are not readily soluble in water. In most cases however, the water and water-ethanol migrates share a large part of features. For instance, the chemical features migrating from a hydration bladder (PUR 2) into water constituted almost an exact subset of those migrating into ethanol (Figure S5 L). These findings underscore the variations in migration patterns and chemical composition across different plastic FCAs.

#### Identification of plastic chemicals

In total, 272 compounds, were tentatively identified, corresponding to 253 features (1.4%) in positive and 94 (1.6%) in negative ionization mode, with an overlap of eleven compounds detected in both (Table S17 and 18). This corresponds to 0– 10% identified features per sample (Table S11). The identification rate is lower than in our previous work,^15,29^ where we included spectral data from the Norman database for identification. To focus our identifications on plastic chemicals, here we limited our search to known compounds used or present in plastics derived from the PlastChem database^4^ and the empirical spectra from MassBank. The limited scope of these two sources explains the lower identification rate obtained here and, once more, highlights the large number of unknown plastic chemicals.

The presence of a compound in all three solvents indicates that it migrates readily and becomes available for human and environmental exposures. Of these features detected in all three solvents, we tentatively identified 74 compounds (2%, Table S19), which include ten known hazardous chemicals as identified in the PlastChem report^4^ (Table S12). These include the antioxidants and stabilizers methyl 3-(3,5-di-tert-butyl-4-hydroxyphenyl)propionate (CAS 6386-38-5) and bis(2,2,6,6-tetramethyl-4-piperidyl) sebacate (CAS 52829-07-9), both toxic to reproduction and to the aquatic environment and nonaethylene glycol nonylphenyl ether (CAS 26571-11-9) an EDC with aquatic toxicity. Of the identified compounds, 15 were tested in ToxCast^21^ for PXR, PPARγ, ERα, or Anti-AR activity, with ten compounds activating at least one of the receptors investigated here (Table S13). Among them, octylparaben (CAS 1219-38-1), detected in seven samples and activating all four receptors tested, and lauryldiethanolamine (CAS 1541-67-9), detected in four PP samples and one PS extract and activating all receptors but ERα. These results confirm the presence and leachability of hazardous chemicals from plastic FCAs.

### 3.2 Receptor activity of FCAs

All plastic FCAs leached chemicals that activated one or more nuclear receptors, while the procedural blanks were inactive. In general, methanol extracts induce significantly stronger effects than the migrates (Figure 2, Figure S7). However, all migrates activated at least two of the receptors tested. In most cases, water-ethanol migrates induced more activity compared to the water migrates.

**Figure 2.**
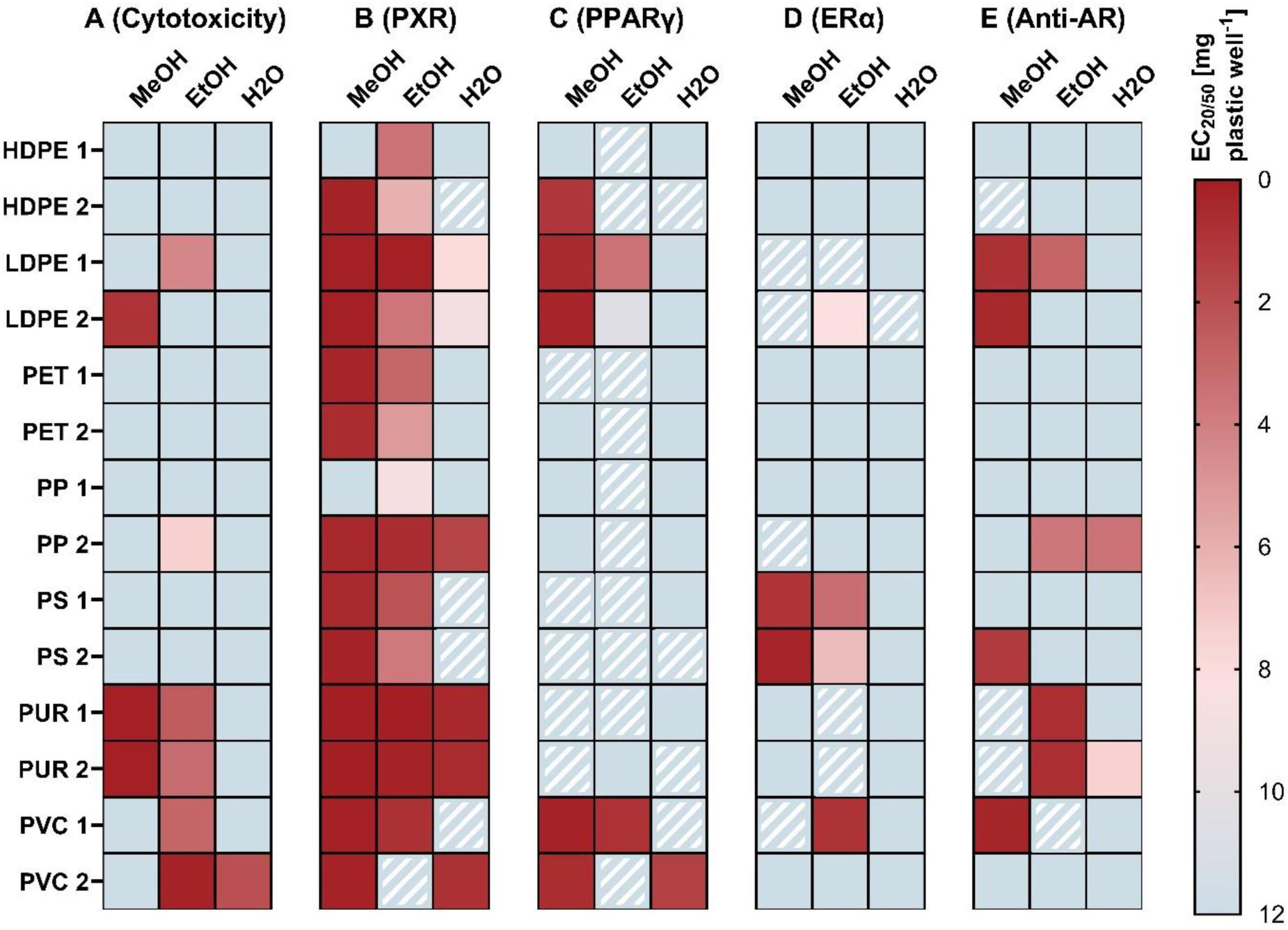
Comparison of toxicity of plastic food contact articles extracted with methanol, migrated into a water-ethanol mixture and water. EC_20_ (EC_50_ for antiandrogenic activity) as mg plastic well^-1^. The highest tested concentration (HTC) was 1.5 mg well^-1^ for the extracts and 12 mg well^-1^ for the migrates, except for cytotoxic samples. If no effect concentration could be derived the value was set to the HTC (12 mg well^-1^, light blue). Stripes = sample was active above the limit of detection, but no EC_20/50_ value could be derived.

#### Cytotoxicity

Cytotoxicity was more prevalent in the water-ethanol migrates as compared to the methanol extracts, mostly due to the higher concentrations tested in the migrates (Figure 2A, Figure S8). A cling film (PVC 2), however, leached cytotoxic chemicals into water-ethanol at concentrations that were non-toxic with the methanol extract (water-ethanol EC_20_ = 0.1 mg well^-1^). Moreover, cytotoxic chemicals from the cling film (PVC 2) also migrated into water.

Cytotoxicity might have led to a masking of the receptor response as indicated by weak receptor activities detected at the lower, non-cytotoxic concentrations (e.g., water-ethanol migrate of PVC 2). However, cytotoxicity can also induce nonspecific receptor activation, which can lead to an overestimation of receptor activity.^30,31^

#### PXR activity

PXR was the predominant target of the plastic chemicals with all FCAs leaching active chemicals into water-ethanol and ten samples leaching these into water (Figure 2B, Figure S9). The methanol extracts induced a significantly stronger effect (mean EC_20_ = 0.19 mg well^-1^) compared to the migrates (mean water-ethanol EC_20_ = 2.86, mean water EC_20_ = 3.30 mg well^-1^). The PUR hydration bladders induced the strongest PXR activity across all leachates. None of the PXR activity migrated from the PET products, HDPE 1 and PP 1 into water. The prevalence of receptor activity across the extracts and migrates reflects the ability of PXR to bind structurally diverse compounds.^32^ Further, it shows that plastic FCAs not only contain, but also leach PXR agonists, suggesting a potential human and environmental exposure to PXR activating chemicals. While PXR is commonly considered a xenobiotic sensor, its role extends beyond this function, encompassing the regulation of energy homeostasis, including glucose, lipid and bile acid metabolism.^33–35^ Moreover, drug-induced dysregulation of PXR is associated with adverse health effects, such as hypercholesterolemia and cardiovascular diseases.^36,37^ Notably, exposure to the plastic chemical dicyclohexyl phthalate has also been linked to PXR-induced atherosclerosis in mice.^38^

Interestingly, the chemicals leaching from two products, a freezer bag (LDPE 1) and a hydration bladder (PUR 1), induced non-monotonic dose responses with a pronounced decrease of PXR activity at higher, non-cytotoxic concentrations (Figure S9, K and S10). Non-monotonic responses are well-described for many EDCs, including plastic chemicals such as BPA.^39,40^ Various mechanisms can contribute to non-monotonic effects mediated through nuclear receptors. Such effects may arise from either individual compounds or compound mixtures.^41^ Given the large number of chemicals detected in these samples, mixture effects are likely. In mixtures, factors such as differences in potency of the individual compounds or response modulations via multiple binding sites within a receptor can play a role.^41,42^

#### PPARγ activity

PPARγ agonists were extracted or migrated from all FCAs (Figure 2C, Figure S11). However, PPARγ activity was mostly lower than 20% compared to the reference compound and only eight samples induced effects >20%. The methanol extracts induced the strongest effects (five samples >20%). However, all but one water-ethanol migrate and five water migrates activated PPARγ, with three and one sample inducing effects >20%, respectively. The most pronounced effects were detected in the PE and PVC products. These results indicate that low concentrations or weak PPARγ agonists migrate from many plastic FCAs. PPARγ, recognized as the master regulator of adipogenesis,^43^ is linked to obesity and metabolic syndromes.^44^ Moreover, plastic extracts have been shown to induce adipogenesis in murine preadipocytes.^45^ However, it remains to be investigated whether the receptor activation observed here can induce effects on cellular or organismal levels.

#### Estrogenic activity

Estrogenic activity was detected in six FCAs with a stronger mean potency found in methanol extracts (Figure 2D, Figure S12). However, the water-ethanol migrate of a freezer bag (LDPE 1, no EC_20_) and the PVC tube (PVC 1, EC_20_ = 0.85 mg well^-1^) induced stronger estrogenicity compared to the corresponding methanol extracts (no EC_20_, Figure S12C and M). Estrogenic chemicals from six FCAs made of LDPE, PS, PUR and PVC migrated into the water-ethanol mixture, whereas migration into water was detected for one freezer bag (LDPE 2). In line with previous research findings^46^, none of the HDPE and PET samples activated the ERα. These results suggest a limited migration of estrogenic chemicals present in PS, PVC and LDPE packaging into water, consistent with findings from another study.^29^ In our previous study,^15^ we detected a significant stronger estrogenicity in PS FCAs as compared to other polymers. Interestingly both PS products tested here leached estrogenic compounds into water-ethanol but not into water. This difference in the estrogenic profile based on the solvents is reflected in the chemical composition, which differed notably between the more lipophilic solvents and water (Figure S5I and J). Considering the capacity of estrogenic compounds to interfere with the endocrine system, leading to adverse developmental and reproductive effects,^47^ these findings underscore the importance of further investigating the causative compounds and potential downstream effects.

#### Antiandrogenic activity

We detected antiandrogenicity in seven FCAs and found that in most cases the activity also migrated into water-ethanol mixtures (Figure 2E, Figure S13). Moreover, antiandrogenic chemicals from a coffee cup and a hydration bladder (PP 2 and PUR 2) also leached into water (mean EC_50_ = 5.4 mg well^-1^). HDPE, PET, PS, and PVC articles did not leach antiandrogenic chemicals into water or water-ethanol. While no EC_50_ was obtained for the PUR extracts, these were much more potent than the corresponding migrates (Figure S13K and L). A decrease in cell number, ranging from 4–12% at the highest tested concentration of the cytotoxic samples, can result in false-positive results in bioassays run in antagonist mode.^48^ To mitigate this, we normalized receptor activity to the cell number in each replicate. Additionally, for most samples we observed antagonistic responses at concentrations that were not cytotoxic at all (Figure S14). The observed rate of antiandrogenic activity migrating into water was comparatively lower than that reported by Zimmermann et al.^29^, possibly due to the higher concentrations of plastic tested there. In another study ^49^, the migration of antiandrogenic cyclic polyester oligomers from PUR adhesives in food packaging was documented.

#### General observations

Consistent with our results on the toxicity of plastic extracts^15,46^ the chemicals migrating from products made of PVC, PUR, and LDPE induced the highest toxicity, whereas PET and HDPE migrates were less toxic. This highlights the need to consider the suitability of specific polymer types for the use as food contact materials. With a few exceptions, most of the tested FCAs leached endocrine and metabolism-disrupting activity into food simulants. As anticipated, the water-ethanol mixture exhibited a higher frequency and potency of receptor activity compared to water. However, it is worth noting that all samples, except the hydration bladders, are designed for use with foods containing higher lipid contents. Therefore, the results for water-ethanol migrates are most relevant here. Notably, the two PUR drinking bladders, intended solely for water use, were found to release endocrine and metabolism disrupting chemicals into the water. Taken together, these results put into question the suitability and safety of these materials for use as FCAs.

### 3.3 Prioritization of chemicals

We followed two strategies to narrow down the chemical complexity of plastic food packaging, first using different solvents for extraction and migration, and secondly applying a multivariate statistical approach using PLS regression. In the first approach, we compared the toxicity profiles and the chemical composition of the three samples derived from the same FCA. By assuming that the toxicity of a product stems from a shared set of chemicals present in the active samples, our focus was on identifying common features among these active samples. For this purpose, we selected three sets of FCAs exhibiting suitable toxicity and chemical profiles: both PUR hydration bladders (PUR 1 and 2) exhibiting PXR and antiandrogenic activity, the PP coffee cup (PP 2) also inducing PXR and antiandrogenic activity, and both PS products which have a pronounced estrogenic activity in the methanol extracts and water-ethanol migrates.

All six PUR samples activate the PXR and are antiandrogenic (tendency for PUR1 water), suggesting that active chemicals might be among the fraction of shared features. We detected 155 shared features across all six active samples, representing 1% of the 13 960 features derived from the hydration bladders (Figure 3A). These shared features serve as a promising starting point for further investigations into compounds driving PXR activity and antiandrogenic effects. Given the pattern of decreasing toxicity from methanol to water-ethanol and water, we refined the analysis by selecting features exhibiting decreasing abundance (as a proxy for concentration) across these solvents, resulting in 33 candidates. Among these we tentatively identified two compounds, 9,10-dihydroxy-hexadecanoic acid (CAS 29242-09-9) and (1-hydroxy-2,4,4- trimethylpentan-3-yl) 2-methylpropanoate (CAS 74367-33-2) with neither having data on receptor activity nor use available.^21,50^ Of the 155 shared features, four additional compounds were tentatively identified, but none was tested for PXR nor antiandrogenic activity (Table S14). However, the antioxidant tris(5-tert-butyl-4-hydroxy-o-tolyl)butane (CAS 35641-51-1) was predicted to have antiandrogenic activity.^51^

**Figure 3.**
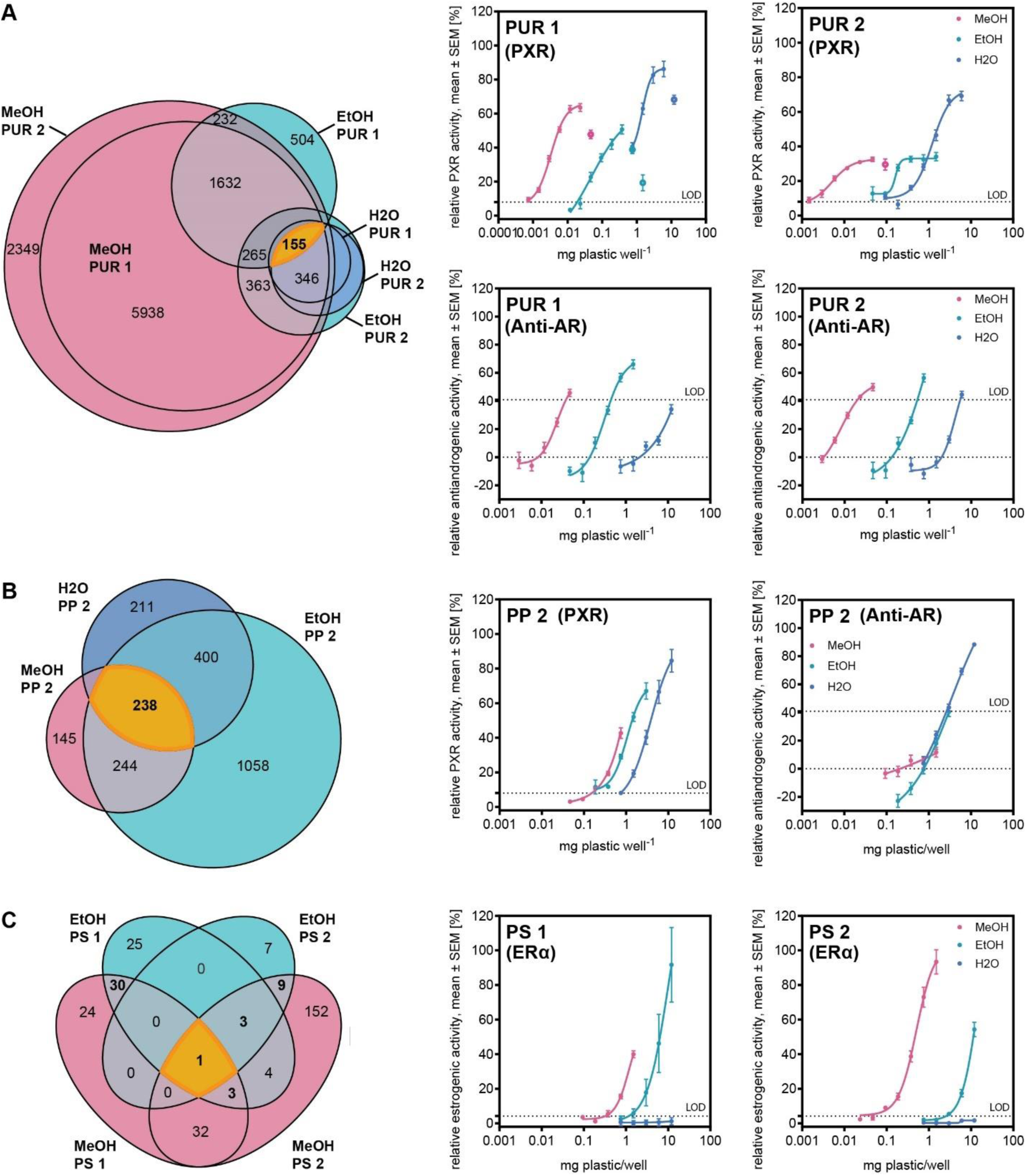
Comparison chemical composition and toxicity migrating to methanol, water-ethanol and water for selected samples: PUR (A), PP 2 (B) and PS (C). Data for dose-response relationships are derived from at least three independent experiments, with four technical replicates per concentration (n ≥ 12).

The three samples derived from the PP coffee cup (PP 2) induced PXR activity with similar potencies in methanol and the water-ethanol mixture, and less activity in the water migrate. A total of 238 features were shared between the three samples potentially containing active chemicals (10% of all PP2 features, Figure 3B). Subsequently, we filtered for features with lower abundance in water than in water-ethanol or methanol – mirroring to the toxicity profiles – resulting in a further reduction to 159 features. For antiandrogenicity, where dose-response relationships of the migrates overlapped (methanol extracts were tested at lower, non-active concentrations), we filtered for features present in the water and water-ethanol migrates at similar abundances (± 25%), which resulted in 37 features with the same abundance and toxicity pattern. Of the 238 features, we tentatively identified ten compounds (Table S14). None was tested for PXR activity and only one plasticizer, ethylene glycol monostearate (CAS 111-60-4), was analyzed for antiandrogenic activity.^21^ However, it was not found to be active.

Regarding the PS products, a single chemical feature was consistently detected across all four active samples, yet it was not identified (Figure 3C). This feature, eluting at 9.6 min, has a m/z of 205.1017 with a charge of +1 and is a promising candidate for identification. Since it is possible that estrogenicity in these distinct samples is induced by different chemicals, we examined the FCAs separately and detected 34 features shared between the methanol extract and water-ethanol migrate of the PS bowl (PS 1), and 13 features in the extract and water-ethanol migrate of the expanded PS food tray (PS 2) serving as candidate features for further investigations into their estrogenic activity. Of these features potentially linked to estrogenic activity, two were identified and tested for estrogenic activity, both inactive (Table S14). However, the emulsifier 1- monolaurin (CAS 142-18-7), which is present in both estrogenic samples of the PS bowl (PS 1), has antiestrogenic activity according to ToxCast^21^, indicating this compound binds to the ERα.

One limitation of this approach is that it presumes that the same active chemical leach into all solvents. Accordingly, such approach is unsuitable if the activity is cause by a compound that would exclusively leach into one specific solvent and another one that leaches into a different solvent. However, the likelihood of such scenario appears to be low given that we compare chemicals originating from the same product. Moreover, the presence of antagonistic chemicals in the samples may confound the results and might lead to an exclusion of active chemicals. Further, this approach is restricted to samples with a suitable toxicity and chemical profile. To mitigate some of these limitations and include all FCAs in the analysis, we applied multivariate statistics, specifically PLS regression.

#### PLS regression

We applied PLS regressions across all four receptors, using a stepwise feature reduction strategy based on variable importance on projection to exclude features not contributing to receptor activity. After five to eleven iterations, all four models improved notably (Table 2, S15). The results were validated by running 500 iterations of random feature sets, each containing the same number of features as the optimized models (Table 2). The complexity of the optimized models was reduced for all receptors, encompassing 1% (115 features for ERα) to 30% (5140 features for PXR) of the initial features (Figure 4, Table 2). These features covary with receptor activity, indicating their potential relevance to the toxicity of the plastic samples. However, a causative relationship between the presence of these features and receptor activity must be confirmed experimentally.

**Figure 4.**
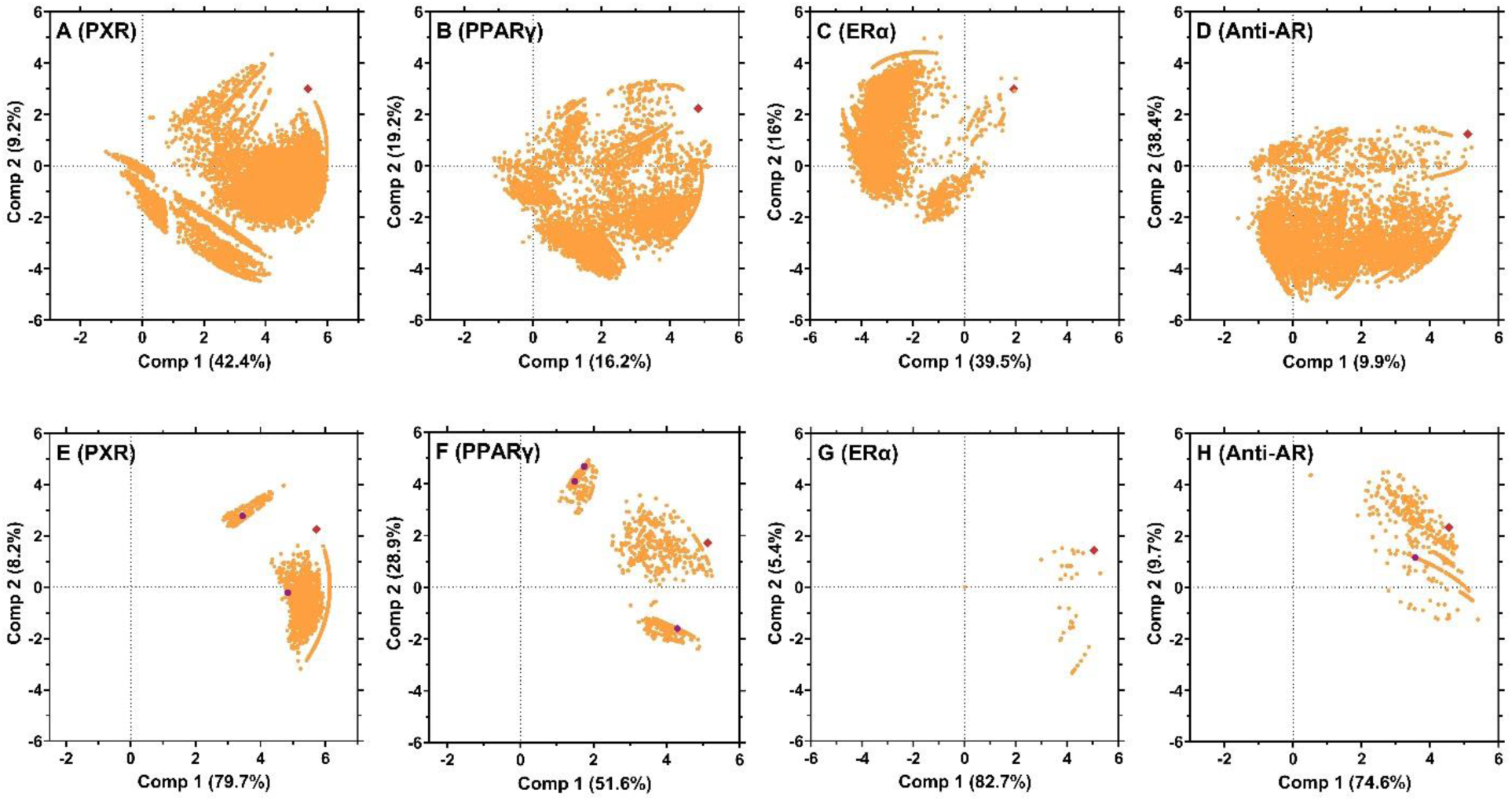
Prioritization of chemical features covarying with the receptor activity using stepwise PLS regressions. Model 0 (A−D) and the optimized models (E−H) for the respective receptors. The receptor activity is represented by red diamonds, chemical features as circles, and tentatively identified features with receptor activity in purple.

**Table 2.** Performance of the PLS regression of chemical features (abundance) and receptor activity (normalized EC_20/50_). Comparison of cross-validated (CV) root mean squared errors (RMSE) and CV determination coefficient (R^2^) of the initial model 0 and optimized model (VIP filtered). Number of features included in the optimized model and corresponding values for the validation based on 500 iterations with randomly selected variables.

| Activity | RMSE CV model 0 | RMSE CV optimized model | R<sup>2</sup> CV optimized model | N features optimized model | PLSR No | RMSE CV random |  | R<sup>2</sup> CV random |  |
| --- | --- | --- | --- | --- | --- | --- | --- | --- | --- |
|  |  |  |  |  |  | min | max | min | max |
| PXR | 84.4 | 78.5 | 0.26 | 5140 (30%) | 9 | 85.9 | 88.4 | 0.06 | 0.11 |
| PPARγ | 1.04 | 0.63 | 0.77 | 2042 (16%) | 8 | 1.2 | 1.3 | 0.07 | 0.23 |
| ERα | 0.92 | 0.75 | 0.24 | 115 (1%) | 5 | 0.9 | 1.3 | -1.24 | 0.01 |
| Anti-AR | 0.93 | 0.55 | 0.58 | 909 (6%) | 11 | 0.8 | 1.0 | -0.52 | 0.00 |

In line with our previous study^15^, the stepwise PLS regression with PXR resulted in the least reduction of features, indicating the wide range of compounds potentially interacting with this receptor. On the contrary, the model for estrogenic activity led to a pronounced reduction of features, resulting in 115 features (1%) after five iterations. One feature in this model was tentatively identified as heptyl decanoate (CAS 60160-17-0), for which no toxicity or usage information was available in PubChem or ToxCast.^21,50^ Nevertheless, this set of features can serve as a promising starting point for further investigations. Both PLS regression models for PPARγ and antiandrogenic activity improved considerably through the feature reduction strategy, comprising 6% (Anti-AR) to 16% (PPARγ) of the initially included features. Interestingly, both models shared 785 features, constituting 86% of the features in the optimized model for antiandrogenicity. This aligns with our previous study,^15^ indicating a correlation between antiandrogenic and PPARγ activity of FCA extracts, suggesting that similar chemicals present in plastics drive both receptor activities.

Of the chemical features which were prioritized using the stepwise PLS regressions, 62 unique compounds were tentatively identified, comprising 0.4% (PXR) to 2% (PPARγ) of the features covarying with receptor activity in the optimized models (Table S20). Of these 62 compounds only nine were tested in ToxCast^21^ for PXR, PPARγ, or antiandrogenic activity. Notably, the plasticizer and flame retardant triphenyl phosphate (CAS 115-86-6) was present in the optimized models of PXR and Anti-AR and is known to activate both receptors (Table S16). Indeed, triphenyl phosphate was recently proposed by the European Chemicals Agency for identification as a substance of very high concern due to its endocrine disrupting properties.^52^ In the PPARγ model, three tentatively identified compounds have PPARγ activity, the most potent being 2-pentylfuran (CAS 3777-69-3), a fragrance and flavoring agent (Table S16).^21,50^ Interestingly, octylparaben, which was tentatively identified in all three solvents and activates all four receptors, was also prioritized in the PXR and PPARγ model.

Our findings underscore the potential of PLS regression with feature reduction based on VIP in reducing the complexity of the chemical landscape of plastics and prioritizing chemical features potentially causing the observed toxicity. However, this approach depends, among others, on the profile of the sample included in the analysis and works better for receptors with more specific binding sites, as indicated by the results for ERα. For the other receptors, the stepwise PLS regressions returned too many features and an additional approach to data reduction would be required.

### 3.5 Implications

Our results confirm that many plastic FCA leach compounds that interfere with nuclear receptors involved in the endocrine and metabolic system and associated with adverse health effects. The chemicals migrating from food packaging made of PVC, PUR, and LDPE induced most endocrine and metabolism disrupting effects, whereas PET and HDPE contained and released less toxicity. While fewer chemicals migrated into water compared to water-ethanol food simulants, several water migrates contained receptor (ant)agonists. This shows that plastics are a relevant source of exposure to toxic chemicals. The approaches explored here for selecting candidate features result in a significant reduction in complexity. However, the limitations in identifying and assessing the toxicity of compounds highlight the need for a more thorough understanding of the chemicals used or present in plastics. This knowledge is essential for improving the identification of toxic plastic chemicals and ultimately creating safer plastics.

## Acknowledgments

This project has received funding from the European Union’s Horizon 2020 research and innovation program under grant agreement No 860720. Special thanks to Andrea Faltynkova for her valuable help in developing the PLS regression models, and to Dru Jagger, Jaeho Lee, and Mara McPartland for their help with obtaining our samples.

## Author contributions

S.S., J.V., and M.W. conceived the study, S.S. performed the sample preparation and the bioassay experiments, Z.B. performed the chemical analyses, S.S. and M.W. analyzed the data, S.S. wrote the manuscript, and all authors provided comments on the manuscript.

## Supporting Information

Supplementary data to this article can be found online.

