## Supporting Information PDF for "Migration of endocrine and metabolism disrupting chemicals from plastic food packaging"

**This file includes:**

S1 Supporting methods and materials

S2 Supporting results and discussion

Figures S1 to S14

Tables S1 to S16

Supplementary references

Other supplementary materials for this manuscript include the following:

Table S17-S20

Access to mass spectral data for all samples is available at https://doi.org/10.18710/QMMER5.

#### S1 Supporting Methods

Table S1. Sample names corresponding to the ones used in our previous study.[1]

| polymer | sample name | sample name in previous study |
|---|---|---|
| HDPE | HDPE 1 | HDPE 2 |
| | HDPE 2 | HDPE 3 |
| LDPE | LDPE 1 | LDPE 1 |
| | LDPE 2 | LDPE 5 |
| PET | PET 1 | PET 1 |
| | PET 2 | PET 6a |
| PP | PP 1 | PP 3 |
| | PP 2 | PP 4 |
| PS | PS 1 | PS 3 |
| | PS 2 | PS 5a |
| PUR | PUR 1 | PUR 1 |
| | PUR 2 | PUR 2 |
| PVC | PVC 1 | PVC 1 |
| | PVC 2 | PVC 3 |

##### S 1.1 Chemical analysis

For chromatographic separation a 20-min gradient program was used (Table S2) and compounds eluting between 1 and 18 min were included in the analysis. The flow rate was set to 0.2 mL min-1 and the column temperature was maintained at 45°C. The injection volume was 1 uL. The collision energy ramped from 15 to 45 V, and the scan time was 0.3 s (details in Table S3). The analysis included extracts and migrates from all 14 food contact articles, alongside 2-4 procedural blanks (PBs) for each solvent, and the methanol solvent itself. Five quality controls comprised of pooled aliquots of all samples, along with additional quality controls comprising all methanol extracts, water-ethanol or water migrates (three each), were included in the analysis. Furthermore, quality controls containing all PUR and PVC, or PP, PE, PET, and PS per solvent, were analyzed.

For data treatment in the software Progenesis QI (Nonlinear Dynamics, version 3.0), the retention times were automatically aligned by the software. Peak picking settings were left in default sensitivity. The minimum chromatographic peak width was set to 0.05 min and the fragment sensitivity was set to 0.2% of the base peak. The program searched for common adducts (positive mode: $[M+H]^+$, $[2M+H]^+$, $[M+2H^{]2+,}$ $[M+H-H_2O]^+$, $M+NH_4$, $[M+Na]^+$, and $[2M+Na]^+$; negative mode: $[M\text{-}H_2O\text{-}H]^-$, $[M\text{-}H]^-$, $[M\text{-}2H]^{2-}$, M+Hac-H), and deconvoluted the mass spectra. The resulting list of chemical features was exported to Microsoft Excel for Windows (Version 2021-2306).

Euler and Venn diagrams to show chemical features shared between samples were produced with the R package "eulerr" (Figure 1A, 3, S7).[2]

Table S2 Chromatographic parameters for the nontarget chemicals analysis.

| Retention time (min) | Mobile phase A (%, water with 0.1% formic acid) | Mobile phase B (%, methanol with 0.1% formic acid) | Curve |
|---|---|---|---|
| 0 | 80 | 20 | - |
| 0.5 | 80 | 20 | 1 |
| 10 | 6 | 94 | 5 |
| 15 | 0 | 100 | 6 |
| 18 | 0 | 100 | 1 |
| 18.1 | 80 | 20 | 1 |
| 20 | 80 | 20 | 1 |

Table S3. Mass spectrometer parameters for the nontarget chemicals analysis.

| Parameter | Value |
|---|---|
| Capillary (kV) | +2.5/-2.0 (positive/negative mode) |
| Source temperature (°C) | 120 |
| Sampling cone voltage (V) | 30 |
| Source offset voltage (V) | 60 |
| Desolvation temperature (°C) | 350 |
| Cone gas flow (L h-1) | 100 |
| Desolvation gas flow (L h-1) | 800 |
| Nebulizer gas flow (bar) | 6 |

#### SPE recovery

To assess the recovery of chemical features in the SPE, the water and water-ethanol migrates of three selected FCAs (HDPE 2, LDPE1 and PVC 1) and PB 1. were analyzed directly (native) without any further treatment (before SPE), and after SPE, following transfer to DMSO and re-dilution to the migration concentration of 0.15 g/mL. Chromatographic separation of the samples was achieved in a 34-minute linear gradient using the same column as described above, a flow rate of 0.2 ml $min^{-1}$ and a column temperature of 55°C. The sample injection volume was 3 uL (Table S4). Two quality controls, comprised of pooled native samples and pooled samples in DMSO, as well as the solvents DMSO and methanol, were included in the analysis alongside the samples. The mass spectrometer settings are the same as for the other samples but were analyzed only in positive ionization mode. The data analysis in Progenesis QI was conducted separately, following the same procedures as described for the other samples, with the exception that the adduct $M+NH_4$ was not included for deconvolution and peak picking utilized automatic sensitivity to detect "fewer" peaks.

To compare feature recovery in the SPE, a hierarchical cluster analysis using Euclidean distances was conducted and visualized in the heatmap (Figure S4) using the R package "pheatmap".[3]

Table S4. Chromatographic parameters for the nontarget chemicals analysis for the SPE recovery analysis.

| Retention time (min) | Mobile phase A (%, water with 0.1% formic acid) | Mobile phase B (%, methanol with 0.1% formic acid) |
|---|---|---|
| 0 | 80 | 20 |
| 0.5 | 80 | 20 |
| 29 | 5 | 95 |
| 36 | 5 | 95 |
| 36.1 | 0 | 100 |
| 38 | 0 | 100 |

#### S1.2 Identification and ToxCast search

To construct the PlastChem database with *in silico* fragmentations, PubChem Identifier Exchange service[4] was utilized to convert the CIDs of the plastic chemicals (n=12 969) from a previous version of the PlastChem report[5] into chemical structures (n=12 297) and retrieved as sdf file. This sdf file was used in the Metascope algorithm of Progenesis QI for the *in silico* fragmentation search.

Table S5. Selected ToxCast assays used to retrieve information about PXR, PPARγ, ERα Anti-AR activity.

| receptor | gene symbol | assay name | cell name |
|---|---|---|---|
| PXR | NR1I2 | ATG_PXRE_CIS | HepG2 |
| | NR1I2 | ATG_PXR_TRANS | HepG2 |
| | NR1I2 | NVS_NR_hPXR | NA |
| | NR1I2 | TOX21_PXR_agonist | HepG2 |
| PPARγ | PPARG | ATG_PPRE_CIS | HepG2 |
| | PPARG | ATG_PPARg_TRANS | HepG2 |
| | PPARG | NVS_NR_hPPARg | NA |
| | PPARG | OT_PPARg_PPARgSRC1_0480 | HEK293T |
| | PPARG | OT_PPARg_PPARgSRC1_1440 | HEK293T |
| | PPARG | TOX21_PPARg_BLA_Agonist_ratio | HEK293T |
| | PPARG | ATG_hPPARg_XSP1 | HepG2 |
| | PPARG | ATG_hPPARg_XSP2 | HepG2 |
| | PPARG | ERF_NR_binding_hPPARG | NA |
| | PPARG | ERFPL_NR_binding_hPPARG | NA |
| ERs | ESR1 | ATG_ERE_CIS | HepG2 |
| | ESR1 | ATG_ERa_TRANS | HepG2 |
| | ESR1 | NVS_NR_hER | NA |
| | ESR1 | OT_ER_ERaERa_0480 | HEK293T |
| | ESR1 | OT_ER_ERaERa_1440 | HEK293T |
| | ESR1 | OT_ER_ERaERb_0480 | HEK293T |
| | ESR1 | OT_ER_ERaERb_1440 | HEK293T |
| | ESR1 | TOX21_ERa_BLA_Agonist_ratio | HEK293T |
| | ESR1 | TOX21_ERa_LUC_VM7_Agonist | VM7 |
| | ESR1 | ATG_hERa_XSP1 | HepG2 |
| | ESR1 | ATG_hERa_XSP2 | HepG2 |
| | ESR1 | CCTE_Deisenroth_AIME_96WELL_LUC_Active | VM7Luc4E2 |
| | ESR1 | CCTE_Deisenroth_AIME_384WELL_LUC_Shift | VM7Luc4E2 |
| AR | AR | ATG_AR_TRANS | HepG2 |
| | AR | NVS_NR_hAR | NA |
| | AR | OT_AR_ARSRC1_0480 | HEK293T |
| | AR | OT_AR_ARSRC1_0960 | HEK293T |
| | AR | TOX21_AR_BLA_Antagonist_ratio | HEK293T |
| | AR | TOX21_AR_LUC_MDAKB2_Antagonist_10nM_R1881 | MDA-kb2 |
| | AR | TOX21_AR_LUC_MDAKB2_Antagonist_0.5nM_R1881 | MDA-kb2 |
| | AR | ATG_hAR_XSP1 | HepG2 |
| | AR | ATG_hAR_XSP2 | HepG2 |
| | AR | UPITT_HCI_U2OS_AR_TIF2_Nucleoli_Antagonist | U2OS |
| | AR | UPITT_HCI_U2OS_AR_TIF2_Nucleoli_Cytoplasm_Ratio_Antagonist | U2OS |
| | AR | ERF_NR_binding_hAR | NA |

#### S1.3 Reporter gene assays

Table S6. Reference compounds used in the bioassays.

| Receptor | Reference compound | CAS | Supplier | Concentration range [mol $^{L-1}$] |
|---|---|---|---|---|
| PXR | nicardipine hydrochloride | 54527-84-3 | Sigma-Aldrich | 9.4 x $10^{-9}$ - 4.8 x $10^{-6}$ |
| PPARγ | rosiglitazone | 122320-73-4 | Sigma-Aldrich | 3.7 x $10^{-10}$ - 3 x $10^{-6}$ |
| ERα | 17-beta-estradiol | 50-28-2 | Sigma-Aldrich | 1.9 x $10^{-13}$ - 1.0 x $10^{-10}$ |
| AR | flutamide | 13311-84-7 | Sigma-Aldrich | 1 x $10^{-8}$ - 3 x $10^{-5}$ |

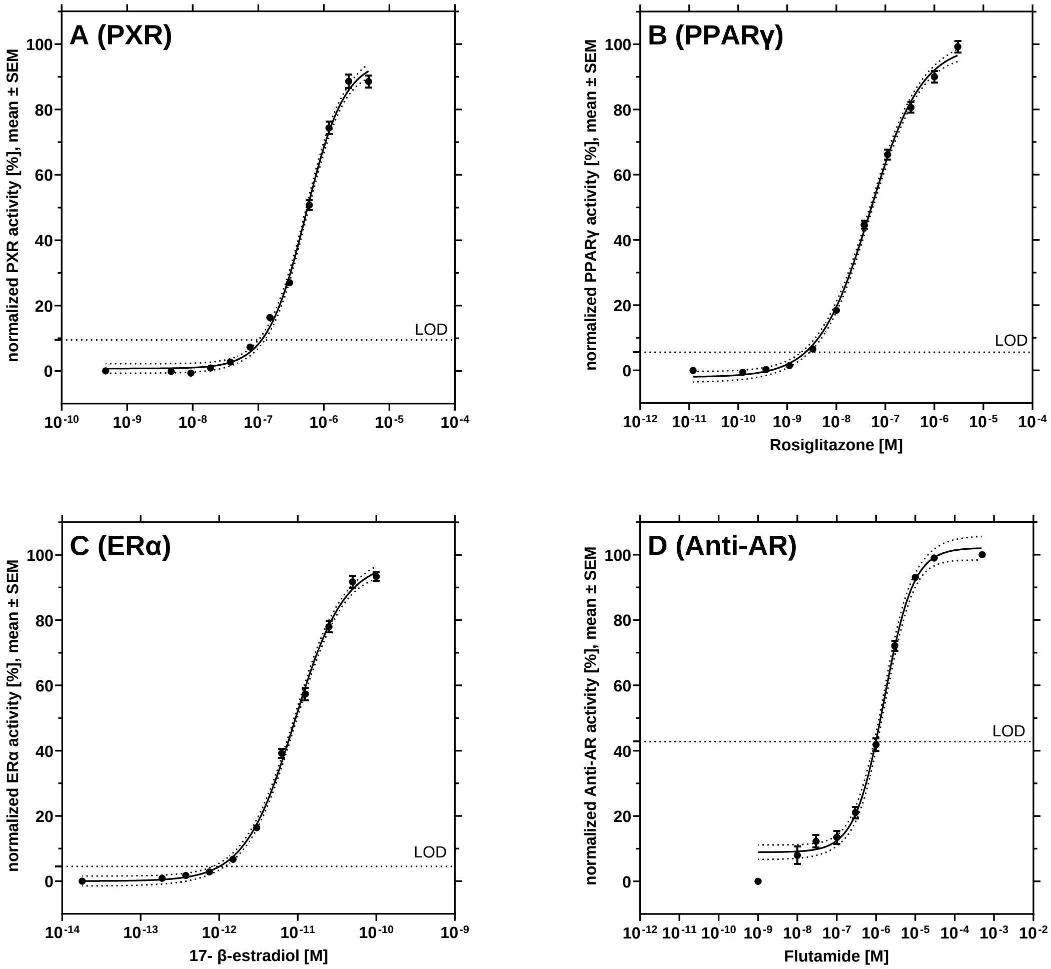


Figure S1. Dose-response relationships of the reference compounds used in the reporter gene assays. Nicardipine was used for the pregnane X receptor (PXR, A), rosiglitazone for peroxisome proliferator receptor gamma (PPARγ, B), 17-β estradiol for estrogen receptor alpha (ERα, C), and flutamide for the anti-androgenic activity (Anti-AR, D). n ≥ 60. Note: LOD = limit of detection

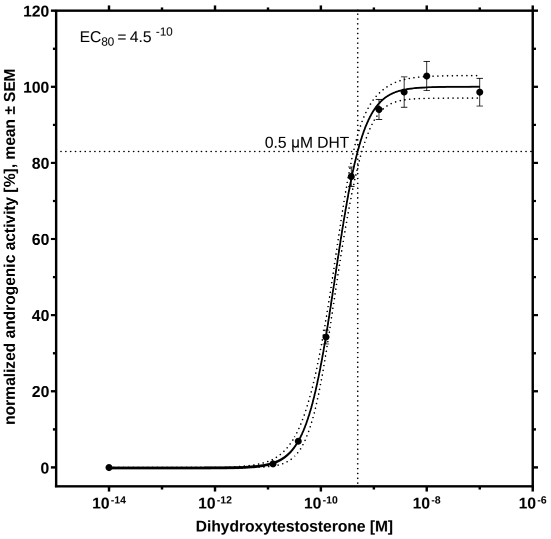


Figure S2. Dose-response relationship of the reference compound dihydrotestosterone (DHT) in the androgen assay. n = 12. Note: 0,5 µM DHT was used as background agonist in the antagonistic assays.

##### S.1.3 PLS regression

Table S7. Number of samples and chemical features included in the initial PLS regression models.

| model | number of samples | number of features |
|---|---|---|
| PXR | 39 | 16956 |
| PPARγ | 31 | 12666 |
| ERα | 32 | 13468 |
| Anti-AR | 30 | 14379 |

### S2. Supporting Results

#### S.2.1 Chemical analysis

##### Recovery of chemical features in the SPE process.

We examined the effect of SPE on the detection of impurities in the procedural blanks (PBs) and found 90-91% of features consistently in the PB before and after SPE, indicating minor losses during SPE (Table S7). Moreover, 9-13% of impurities were added during the SPE, but no additional impact of 10% ethanol was found. However, these impurities were accounted for in the samples, by subtracting the 10-fold abundance of all features detected in the corresponding PBs.

Table S8. Effect of the SPE on the number of chemical features detected in the procedural blanks.

| Procedural blank | features before SPE | recovered in SPE | lost in SPE | added in SPE |
|---|---|---|---|---|
| PB_EtOH | 5141 | 4602 (90%) | 539 (10%) | 480 (9%) |
| PB_H2O | 4995 | 4491 (90%) | 504 (10%) | 661 (13%) |
| PB_H2O 10% EtOH | 4995 | 4524 (91%) | 471 (9%) | 576 12%) |

Table S9. Effect of 10% EtOH on the SPE.

| sample | features before SPE | recovered in SPE | lost in SPE | added in SPE |
|---|---|---|---|---|
| HDPE2_H2O | 81 | 67 (83%) | 14 (17%) | 11 (14%) |
| LDPE1_H2O | 419 | 354 (84%) | 65 (16%) | 97 (23%) |
| PVC1_H2O | 300 | 259 (86%) | 41 (14%) | 32 (11%) |

Table S10. Recovery of chemical features in the SPE.

| sample | features before SPE | recovered in SPE | lost in SPE | added in SPE |
|---|---|---|---|---|
| HDPE 2 EtOH | 1059 | 295 (28%) | 764 (72%) | 49 (5%) |
| HDPE 2 H2O | 72 | 19 (26%) | 53 (74%) | 62 (6%) |
| LDPE 1 EtOH | 3466 | 1904 (55%) | 1562 (45%) | 17 (0.5%) |
| LDPE 1 H2O | 303 | 272 (90%) | 31 (10%) | 147 (49%) |
| PVC 1 EtOH | 2469 | 2023 (82%) | 446 (18%) | 402 (16%) |
| PVC 1 H2O | 186 | 159 (85%) | 27 (15%) | 141 (76%) |

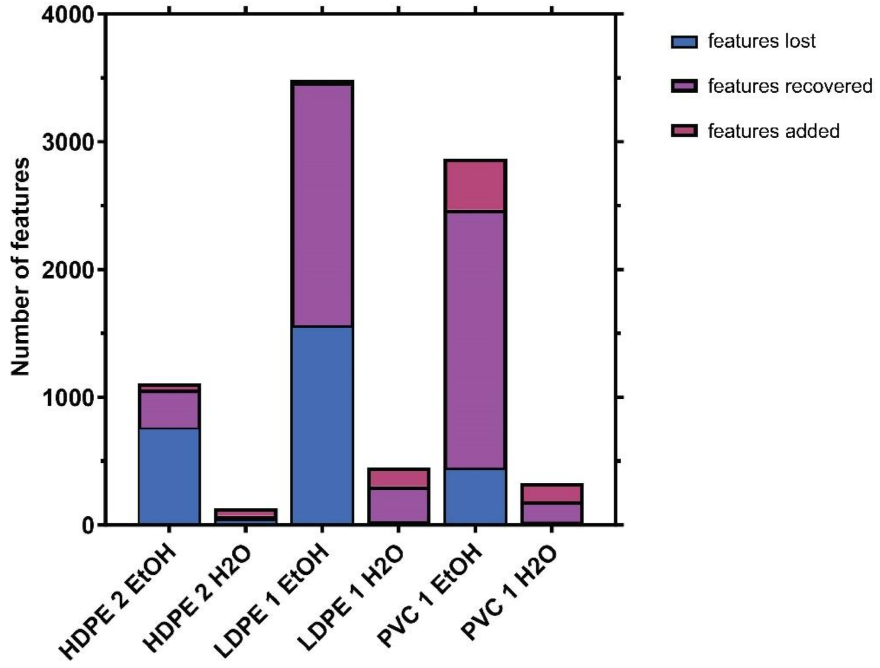


Figure S3. Recovery of chemical features in SPE.

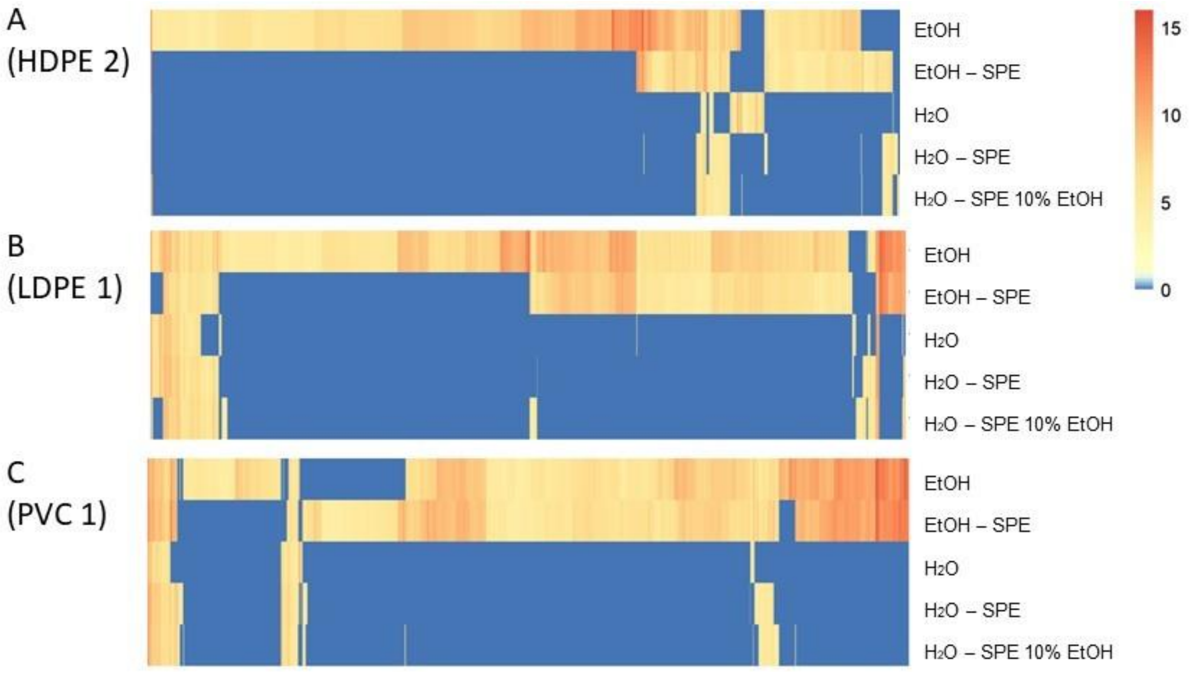


Figure S4. Recovery of chemical features in SPE shown as clustered heatmap.

#### Chemical landscape of the FCAs.

Table S11. Number of features and identified features detected in positive and negative ionization mode.

| Positive ionization | $H_2O$ | | | EtOH | | | MeOH | | |
|---|---|---|---|---|---|---|---|---|---|
| | detected | identified | percent identified | detected | identified | percent identified | detected | identified | percent identified |
| HDPE 1 | 25 | 1 | 4.0 | 64 | 3 | 4.7 | 6 | 0 | 0 |
| HDPE 2 | 8 | 0 | 0 | 35 | 0 | 0.0 | 315 | 5 | 1.6 |
| LDPE 1 | 380 | 11 | 2.9 | 2521 | 85 | 3.4 | 957 | 70 | 7.3 |
| LDPE 2 | 250 | 7 | 2.8 | 1028 | 43 | 4.2 | 1652 | 76 | 4.6 |
| PET 1 | 10 | 1 | 10.0 | 341 | 6 | 1.8 | 253 | 5 | 2.0 |
| PET 2 | 25 | 1 | 4.0 | 110 | 2 | 1.8 | 39 | 2 | 5.1 |
| PP 1 | 29 | 1 | 3.4 | 73 | 1 | 1.4 | 18 | 2 | 11.1 |
| PP 2 | 861 | 23 | 2.7 | 1940 | 31 | 1.6 | 639 | 20 | 3.1 |
| PS 1 | 38 | 2 | 5.3 | 68 | 0 | 0.0 | 90 | 4 | 4.4 |
| PS 2 | 19 | 0 | 0 | 20 | 1 | 5.0 | 202 | 5 | 2.5 |
| PUR 1 | 980 | 25 | 2.6 | 3027 | 36 | 1.2 | 10236 | 107 | 1.0 |
| PUR 2 | 1068 | 29 | 2.7 | 1945 | 40 | 2.1 | 11751 | 105 | 0.9 |
| PVC 1 | 184 | 9 | 4.9 | 2939 | 34 | 1.2 | 2819 | 45 | 1.6 |
| PVC 2 | 3812 | 81 | 2.1 | 10631 | 156 | 1.5 | 6083 | 96 | 1.6 |
| Negative ionization | $H_2O$ | | | EtOH | | | MeOH | | |
| | detected | identified | percent identified | detected | identified | percent identified | detected | identified | percent identified |
| HDPE 1 | 17 | 1 | 5.9 | 58 | 0 | 0 | 23 | 0 | 0 |
| HDPE 2 | 60 | 3 | 5 | 136 | 3 | 2 | 101 | 5 | 5 |
| LDPE 1 | 159 | 5 | 3.1 | 1053 | 15 | 1.4 | 360 | 10 | 2.8 |
| LDPE 2 | 92 | 4 | 4.3 | 344 | 11 | 3.2 | 413 | 10 | 2.4 |
| PET 1 | 37 | 2 | 5.4 | 173 | 7 | 4.0 | 260 | 8 | 3.1 |
| PET 2 | 26 | 0 | 0.0 | 50 | 1 | 2.0 | 22 | 1 | 4.5 |
| PP 1 | 189 | 3 | 1.6 | 57 | 1 | 1.8 | 32 | 0 | 0 |
| PP 2 | 343 | 14 | 4.1 | 304 | 7 | 2.3 | 128 | 3 | 2.3 |
| PS 1 | 29 | 1 | 3.4 | 39 | 2 | 5.1 | 67 | 0 | 0 |
| PS 2 | 79 | 0 | 0 | 47 | 0 | 0 | 87 | 0 | 0 |
| PUR 1 | 428 | 8 | 1.9 | 756 | 7 | 0.9 | 1944 | 35 | 1.8 |
| PUR 2 | 487 | 8 | 1.6 | 516 | 5 | 1.0 | 2388 | 36 | 1.5 |
| PVC 1 | 183 | 13 | 7.1 | 1046 | 17 | 1.6 | 352 | 7 | 2.0 |
| PVC 2 | 2422 | 49 | 2.0 | 3956 | 72 | 1.8 | 1409 | 28 | 2.0 |

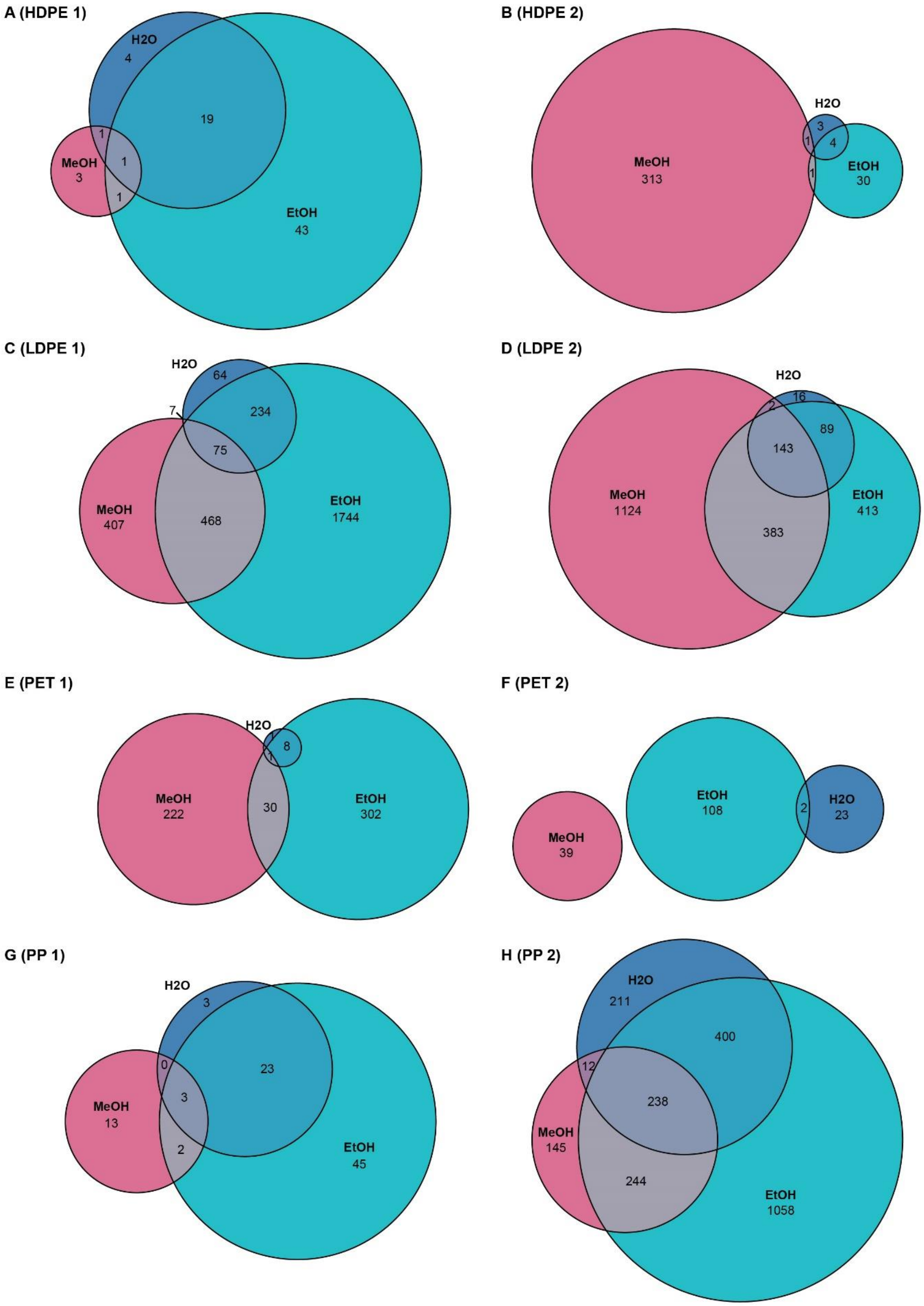
A (HDPE 1)
H2O
4
19
1
MeOH
3
1
1
EtOH
43
B (HDPE 2)
MeOH
313
H2O
3
1
4
1
EtOH
30
C (LDPE 1)
H2O
64
7
234
75
MeOH
407
468
EtOH
1744
D (LDPE 2)
H2O
2
16
89
143
MeOH
1124
383
EtOH
413
E (PET 1)
H2O
1
8
1
MeOH
222
30
EtOH
302
F (PET 2)
MeOH
39
EtOH
108
2
H2O
23
G (PP 1)
H2O
3
0
23
3
MeOH
13
2
EtOH
45
H (PP 2)
H2O
211
400
12
238
MeOH
145
244
EtOH
1058

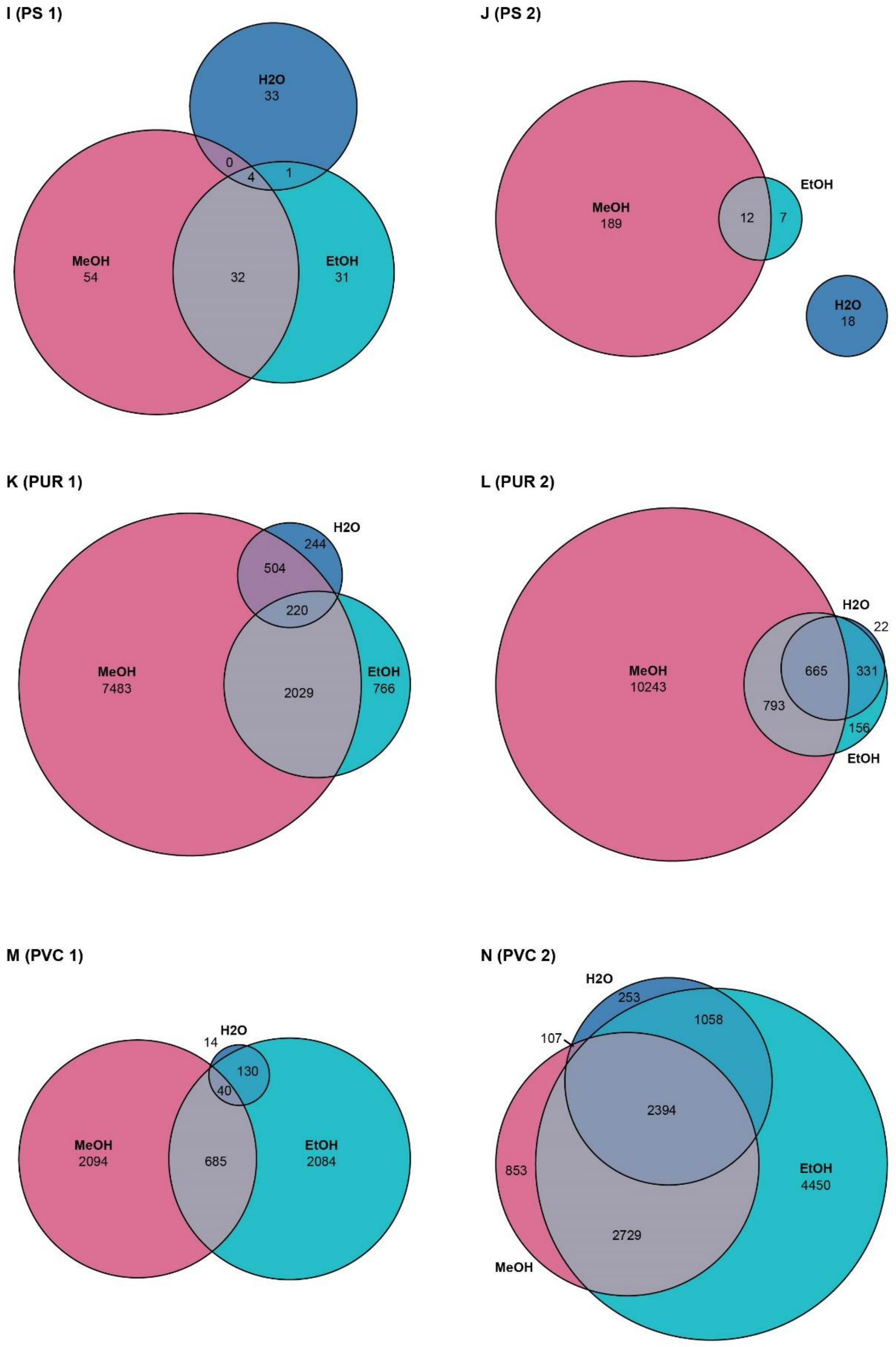


Figure S5. Overlap of chemical features extracted and migrating from one FCA (positive ionization mode). An overlap of <1% is not shown.

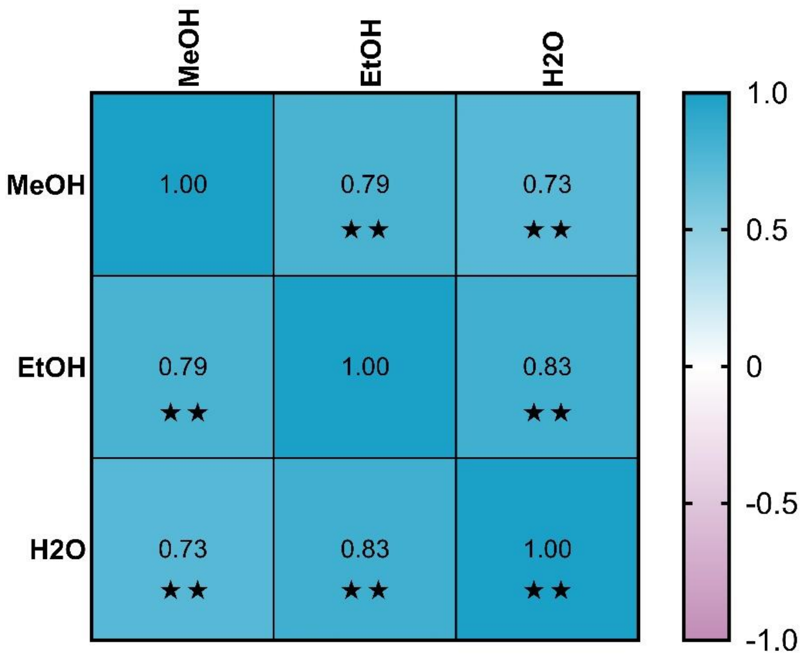


Figure S6. Correlation of the number of features detected with the different solvents.

#### S2.2 Identification of plastic chemicals

Table S12. Tentatively identified chemicals shared across water migrates, water-ethanol migrates and methanol extracts and present in the red list of the PlastChem report[5], along with use and toxicity data. CMR: cancerogenic, mutagenic and toxic to reproduction; STOT: specific target organ toxicity.

| compound ID | compound Name | CAS | CID | toxicity | use |
|---|---|---|---|---|---|
| 10.38_616.4158n | Nonaethylene glycol nonylphenyl ether | 26571-11-9 | 72385 | EDC, aquatic toxicity | Antioxidant; Catalyst; Light stabilizer; Monomer |
| 3.92_255.2198m/z; 4.30_255.2194m/z; 4.97_255.2193m/z; 6.57_255.2198m/z | Bis(1,2,2,6,6-pentamethyl-4-piperidyl) sebacate | 41556-26-7 | 586744 | CMR, aquatic toxicity | Antioxidant; Colorant; Filler; Light stabilizer; Monomer; Other Processing Aids |
| 7.83_273.2667n, 7.83_273.2667n | N-Lauryldiethanolamine | 1541-67-9 | 352309 | aquatic toxicity | |
| 4.24_241.2041m/z, 5.32_480.3927n | Bis(2,2,6,6-tetramethyl-4-piperidyl) sebacate | 52829-07-9 | 164282 | CMR, aquatic toxicity | Antioxidant; Biocide; Colorant; Filler; Heat stabilizer; Intermediate; Light stabilizer; Lubricant; Monomer; Other Processing Aids |
| 6.72_143.0707m/z | Diglycidyl 1,2-cyclohexanedicarboxylate | 5493-45-8 | 21660 | aquatic toxicity | Intermediate; Other Processing Aids |
| 9.34_178.1366n | 3-Methyl-5-phenylpentan-1-ol | 55066-48-3 | 108312 | STOT | Odor Agent |
| 8.88_315.1934m/z | Methyl 3-(3,5-di-tert-butyl-4-hydroxyphenyl)propionate | 6386-38-5 | 62603 | CMR, STOT, aquatic toxicity | Antioxidant; Filler; Light stabilizer; Lubricant |
| 6.06_254.1717m/z | N,N-Bis(2-hydroxyethyl)octanamide | 68155-07-7 | 76499 | CMR, aquatic toxicity | |
| 5.71_370.2947m/z | Methyl 1,2,2,6,6-pentamethyl-4-piperidyl sebacate | 82919-37-7 | 157881 | CMR, aquatic toxicity | Antioxidant; Light stabilizer |
| 7.65_329.1565m/z | 2-Ethylhexyl 10-ethyl-4-((2-((2-ethylhexyl)oxy)-2-oxoethyl)thio)-7-oxo-8-oxa-3,5-dithia-4-phosphatetradecanoate 4-oxide | 83547-95-9 | 93477 | aquatic toxicity | |

Table S13. Tentatively identified chemicals shared across water migrates, water-ethanol migrates and methanol extracts, along with their receptor activity ($AC_{50}$) according to ToxCast.

| **receptor** | **compound name** | **CAS** | **$AC_{50}$ [μM]** | **assay name** |
|---|---|---|---|---|
| PXR | 4-Methylumbelliferone | 90-33-5 | 8.1 | ATG_PXRE_CIS |
| | Octylparaben | 1219-38-1 | 12.2 | ATG_PXRE_CIS |
| | Lauryldiethanolamine | 1541-67-9 | 13.5 | TOX21_PXR_agonist |
| | Bis(2,2,6,6-tetramethyl-4-piperidyl) sebacate | 52829-07-9 | 18.1 | TOX21_PXR_agonist |
| | Dexamethasone | 50-02-2 | 35.6 | TOX21_PXR_agonist |
| | 3-Methyl-5-phenylpentan-1-ol | 55066-48-3 | 43.3 | ATG_PXRE_CIS |
| | Triethylene glycol dimethacrylate | 109-16-0 | 68.6 | ATG_PXRE_CIS |
| | Bis(2,3-epoxypropyl) cyclohexane-1,2-dicarboxylate | 5493-45-8 | 69.0 | TOX21_PXR_agonist |
| PPARγ | Octylparaben | 1219-38-1 | 10 | ATG_hPPARg_XSP2 |
| | Lauryldiethanolamine | 1541-67-9 | 76.93 | TOX21_PPARg_BLA_Agonist_ratio |
| ERα | Octylparaben | 1219-38-1 | 4.7 | TOX21_ERa_LUC_VM7_Agonist |
| | 2-Hydroxyethyl octadecanoate | 111-60-4 | 17.2 | TOX21_ERa_LUC_VM7_Agonist |
| | 4-Methylumbelliferone | 90-33-5 | 42.1 | ATG_ERE_CIS |
| | Dimethoxane | 828-00-2 | 76.8 | ATG_ERE_CIS |
| AR | Dexamethasone | 50-02-2 | 9.3 | UPITT_HCI_U2OS_AR_TIF2_Nucleoli_Antagonist |
| | Lauryldiethanolamine | 1541-67-9 | 17.4 | TOX21_AR_LUC_MDAKB2_Antagonist_10nM_R1881 |
| | Octylparaben | 1219-38-1 | 29.4 | TOX21_AR_BLA_Antagonist_ratio |
| | 3-Methyl-5-phenylpentan-1-ol | 55066-48-3 | 34.2 | TOX21_AR_LUC_MDAKB2_Antagonist_0.5nM_R1881 |

#### S.2.2 Reporter gene assays

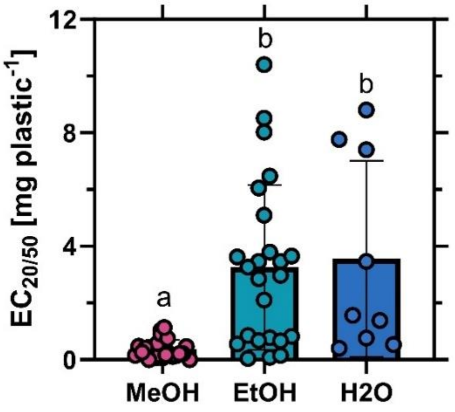


Figure S7. Comparison of receptor activation between solvents. Included are only samples were $EC_{20/50}$ values could be derived. ANOVA with Turkey's multiple comparison tests for statistical differences ($p < 0.05$) indicated by letters. .Note: MeOH extracts were analyzed at lower concentrations (1.5 mg well$^{-1}$) as compared to the migrates (12 mg well$^{-1}$), except for cytotoxic samples.

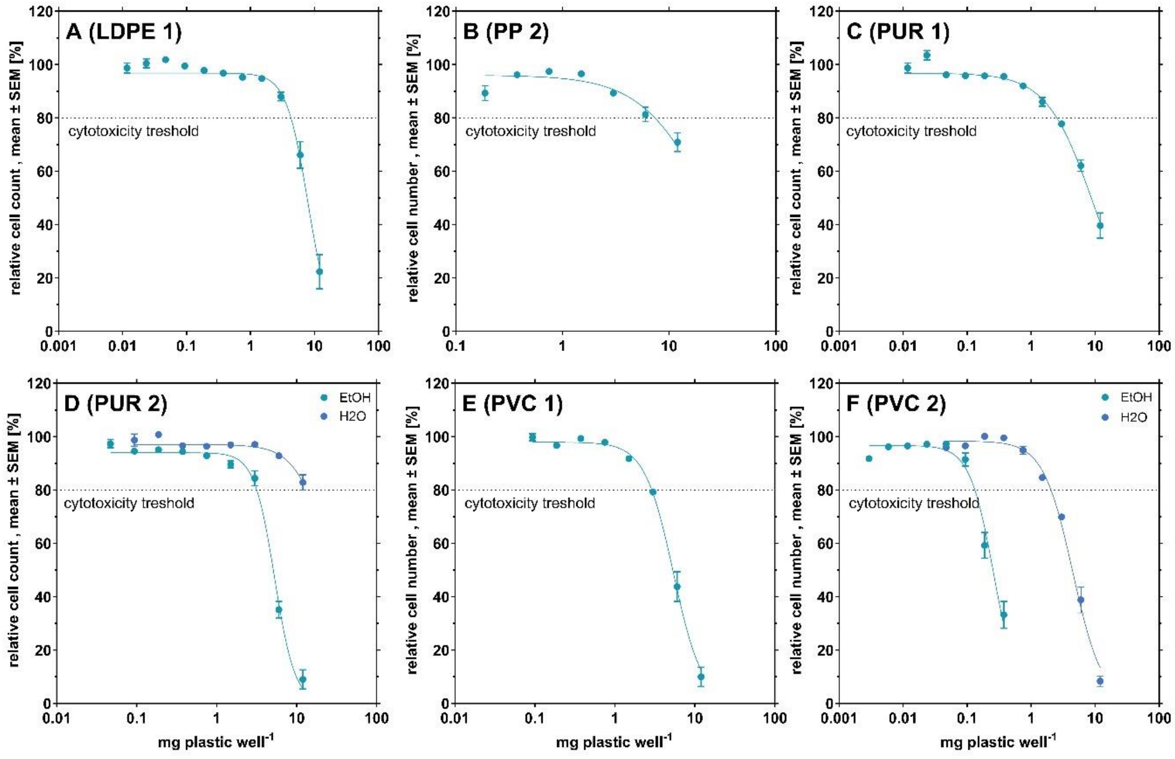


Figure S8. Dose-response relationships of the cytotoxic samples. Cytotoxicity is defined as a cell count <80% of the pooled negative and solvent controls. Cell count across receptors combined, for cytotoxic concentrations $n \geq 16$, for non-cytotoxic concentrations $n \geq 48$. Highest tested concentration of 12 mg plastic/wel

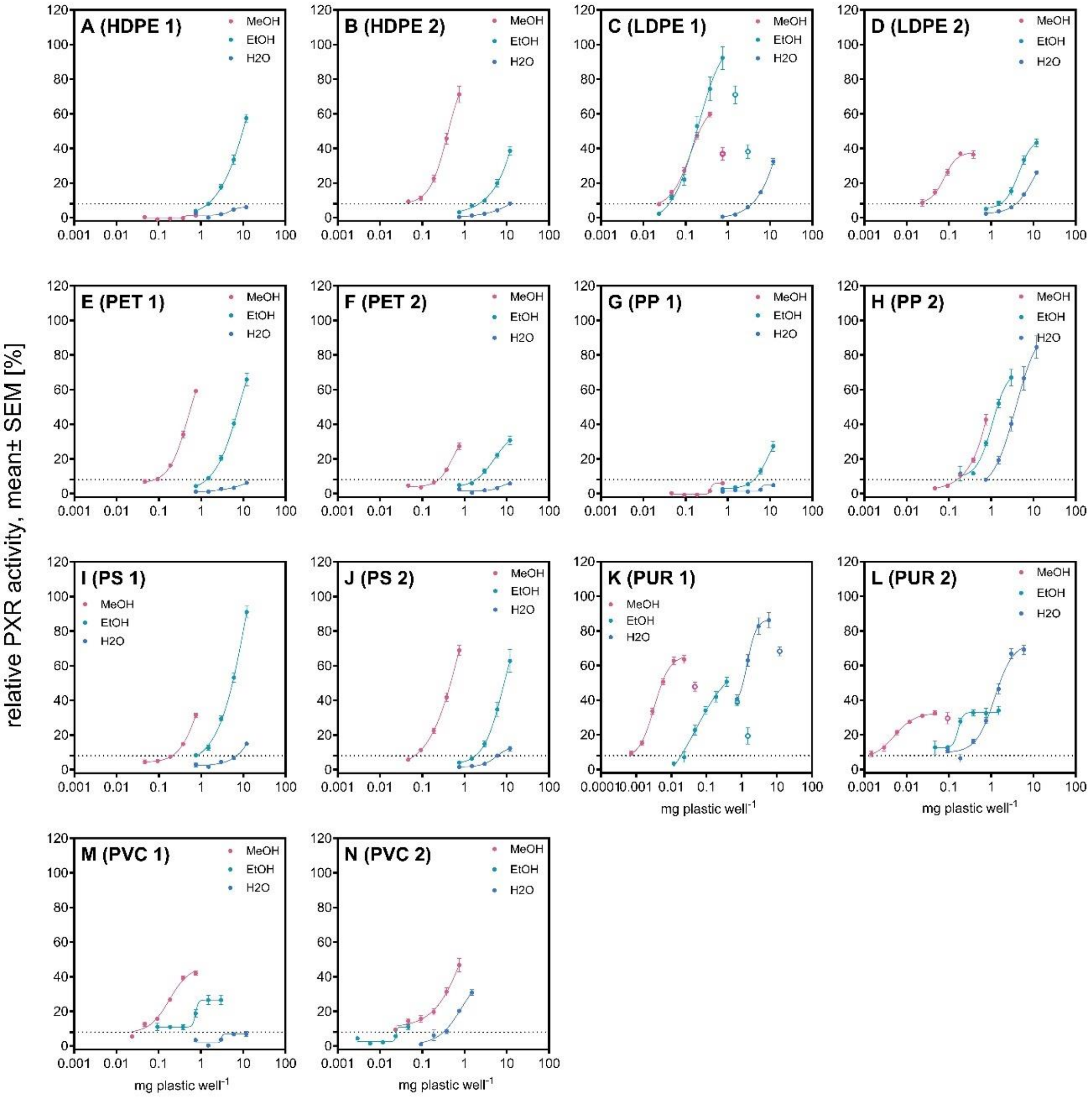


Figure S9. Comparison of PXR activity between methanol extracts, water-ethanol and water migrates. Data are derived from at least three independent experiments, with four technical replicates per concentration (n ≥ 12).

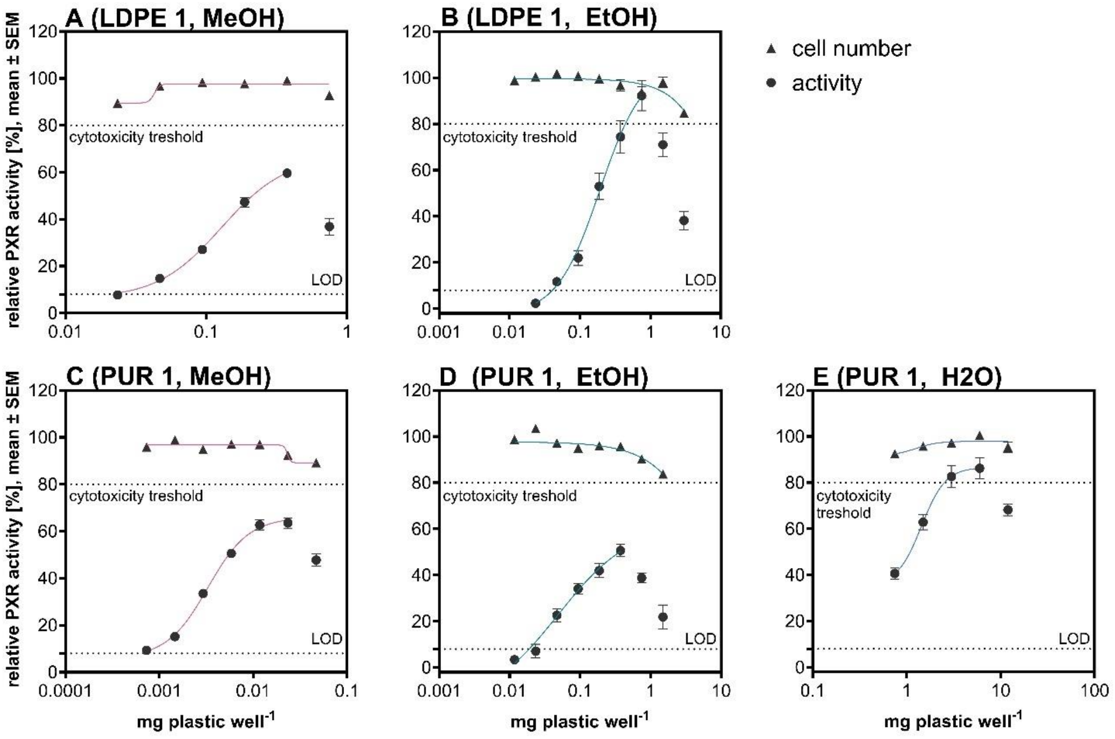


Figure S10. Non-monotonic dose response of PXR activity along with the relative cell number at the tested concentrations. Data are derived from at least three independent experiments, with four technical replicates per concentration (n ≥ 12). LOD = limit of detection.

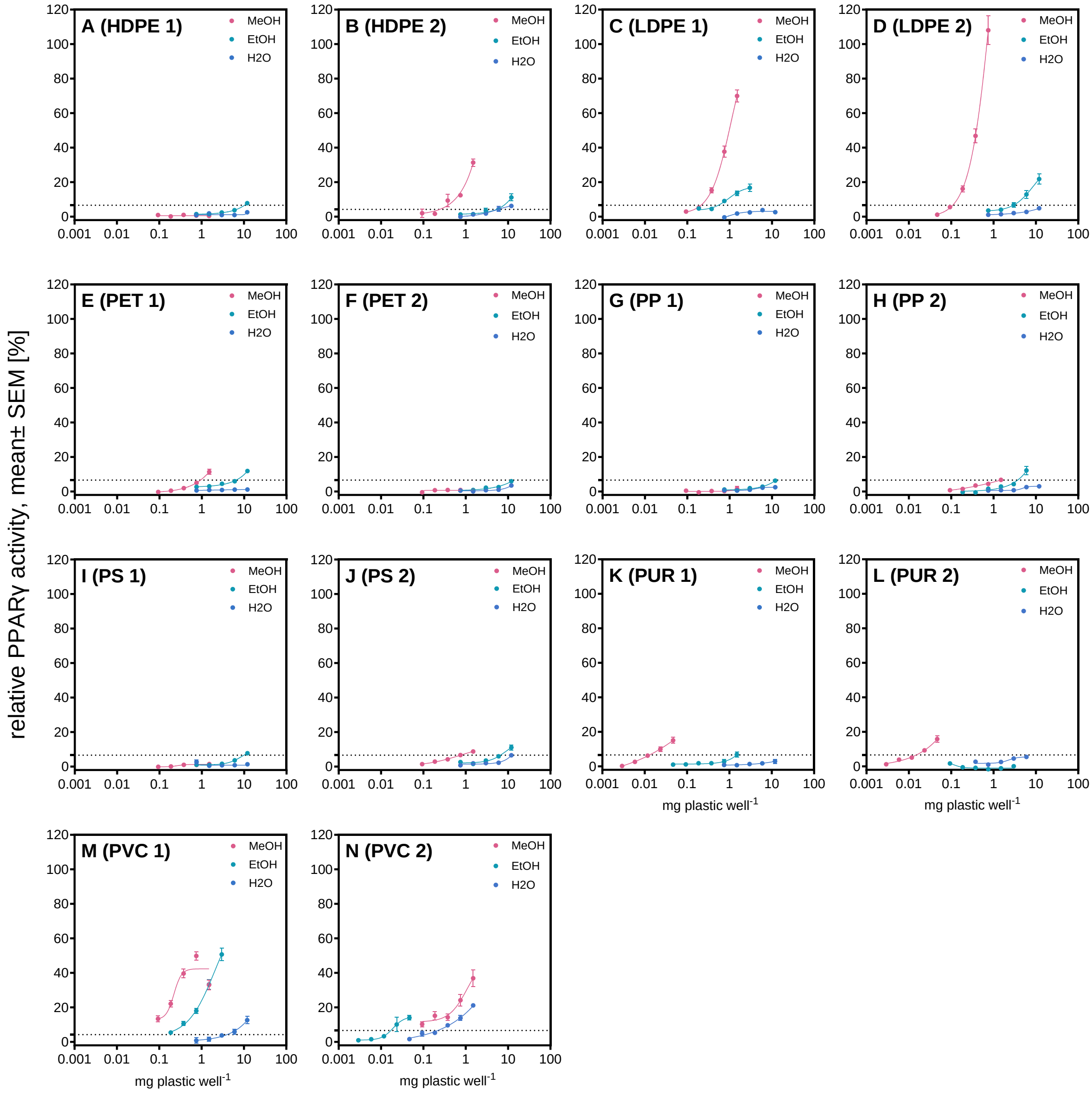


Figure S11 Comparison of PPARγ activity between methanol extracts, water-ethanol (50%) and water migrates. Data are derived from at least three independent experiments, with four technical replicates per concentration ($n \geq 12$).

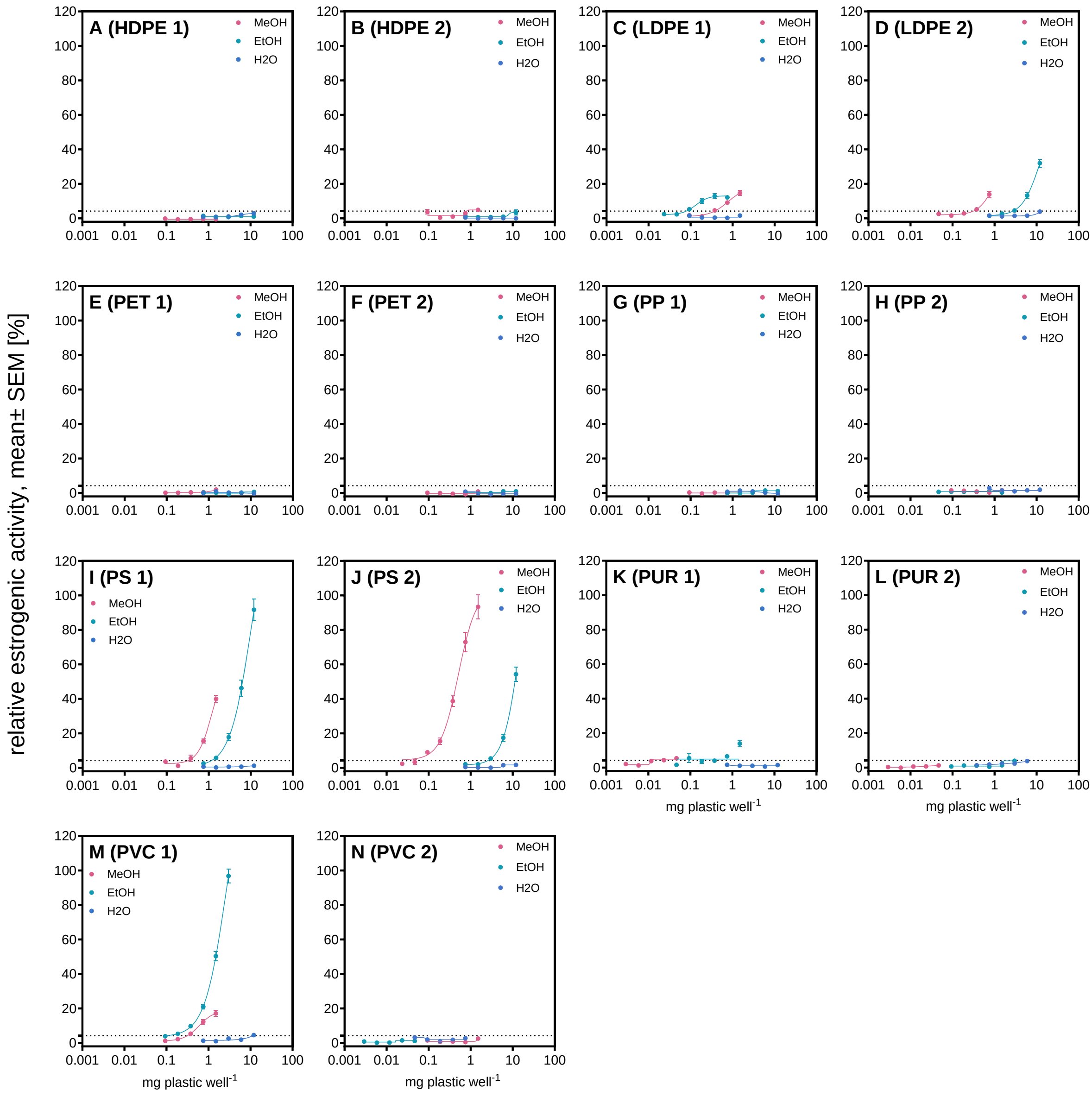


Figure S12. Comparison of estrogenic activity between methanol extracts, water-ethanol and water migrates. Data are derived from at least three independent experiments, with four technical replicates per concentration (n ≥ 12).

.

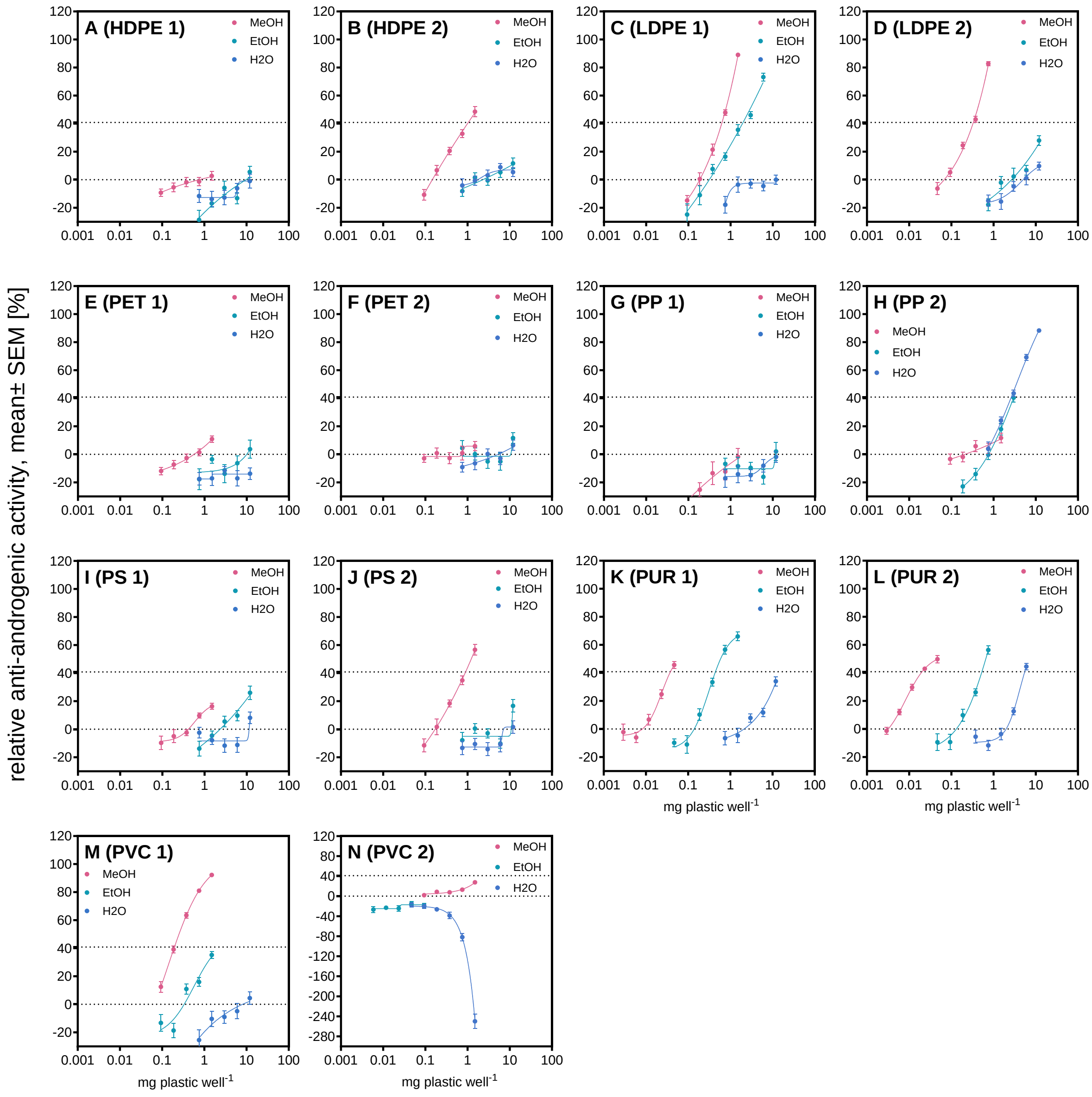


Figure S13. Comparison of anti-androgenic activity between methanol extracts, water-ethanol and water migrates. Data are derived from at least three independent experiments, with four technical replicates per concentration (n ≥ 12).

.

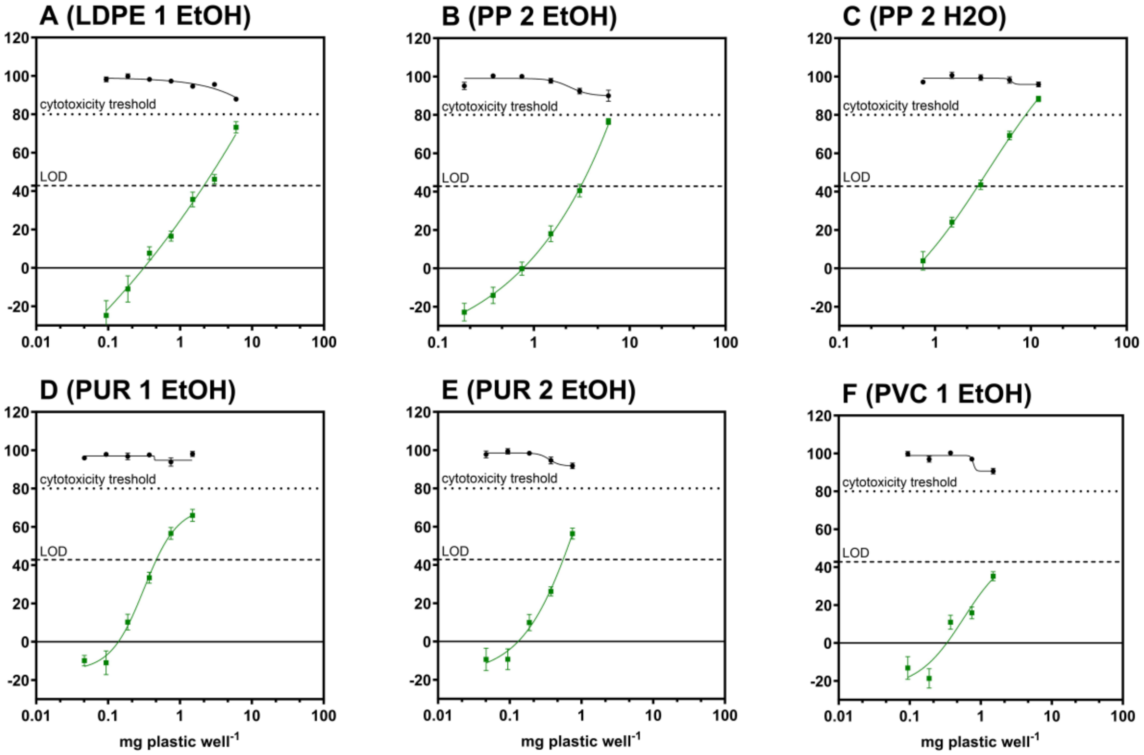


Figure S14. Dose-response relationships of the anti-androgenic migrates with the corresponding cell count (normalized to controls). Data are derived from at least three independent experiments, with four technical replicates per concentration (n ≥ 12).

#### S2.3 Prioritization of chemicals

Table S14. Tentatively identified chemicals selected based on their presence in active samples of the same product/polymer type, along with their receptor activity according to ToxCast.

| FCA | compound name | CID | CAS | PXR | Anti-AR | ERα |
|---|---|---|---|---|---|---|
| **PUR** | Tris(5-tert-butyl-4-hydroxy-o-tolyl)butane | 3015792 | 35641-51-1 | na | na | - |
| | (1-hydroxy-2,4,4-trimethylpentan-3-yl) 2-methylpropanoate | 156477 | 74367-33-2 | na | na | - |
| | 9,10-dihydroxyhexadecanoic acid | 193113 | 29242-09-9 | na | na | - |
| | Adipic acid, methyl vinyl ester | 18084 | 2969-87-1 | na | na | - |
| | Triethylene glycol monomethacrylate | 13908061 | 2351-42-0 | na | na | - |
| | 2-Cyclohexen-1-one, 2-methyl-3-(2-methylpropoxy)- | 317029 | 37457-15-1 | na | na | - |
| **PP 2** | Bis(2,6-di-tert-butyl-4-methylphenyl)pentaerythritol diphosphite | 12868350 | 78837-87-3 | na | na | - |
| | Bis(2,6-di-tert-butyl-4-methylphenyl)pentaerythritol diphosphite | 3601357 | 80693-00-1 | na | na | - |
| | Atractyloside I | 10929902 | - | na | na | - |
| | Triethylene glycol monomethacrylate | 13908061 | 2351-42-0 | na | na | - |
| | 1,6,13,18-Tetraoxacyclotetracosane-7,12,19,24-tetrone | 12868350 | 78837-87-3 | na | na | - |
| | Triethylene glycol monomethacrylate | 13908061 | 2351-42-0 | na | na | - |
| | Ethylene glycol monostearate | 24762 | 111-60-4 | na | inactive | - |
| | 11-ethyl-5-methylpentadecanoic acid | 109214 | 68201-37-6 | na | na | - |
| | (2E,13Z)-2,13-Octadecadienol | 5283306 | 123551-47-3 | na | na | - |
| | Cyclotris(1,4-butylene Terephthalate) | 22186544 | 63440-94-8 | na | na | - |
| **PS 1** | 1-monolaurin | 14871 | 142-18-7 | - | - | inactive |
| | N-Lauryldiethanolamine | 352309 | 1541-67-9 | - | - | inactive |

Table S15. Performance of the PLS regression of chemical features (occurrence and abundance) and receptor activity (normalized EC20/50 values) after iterative filtering steps excluding features with VIP <0.8. Calculated (Cal), cross-validated (CV), cumulative explained variance (cumexpvar) of the chemical features (X) and receptor activity (Y), determination coefficient (R2), root mean squared errors (RMSE), slope for predicted vs. measured values (slope), bias for prediction vs. measured values (bias), ratio of standard deviation of response values to standard error of prediction (RPD).

| Model | N selected components | N features | | X cumexpvar | Y cumexpvar | R2 | RMSE | Slope | Bias | RPD |
|---|---|---|---|---|---|---|---|---|---|---|
| PXR | | | | | | | | | | |
| 0 | 4 | 16956 | Cal | 66 | 100 | 1 | 1.8 | 1 | 0 | 52.2 |
| | | | Cv | NA | NA | 0.14 | 84.4 | 0.42 | -1.8 | 1.1 |
| 1 | 5 | 10684 | Cal | 92 | 100 | 1 | 1.6 | 1 | 0 | 57.2 |
| | | | Cv | NA | NA | 0.19 | 82.1 | 0.43 | -2.2 | 1.1 |
| 2 | 5 | 8288 | Cal | 93 | 100 | 1 | 1.9 | 1 | 0 | 49.0 |
| | | | Cv | NA | NA | 0.20 | 81.8 | 0.45 | -2.2 | 1.1 |
| 3 | 1 | 7493 | Cal | 73 | 81 | 0.81 | 40.1 | 0.81 | 0 | 2.3 |
| | | | Cv | NA | NA | 0.14 | 84.5 | 0.47 | 1.4 | 1.1 |

| Model | N selected components | N features | | X cumexpvar | Y cumexpvar | R2 | RMSE | Slope | Bias | RPD |
|---|---|---|---|---|---|---|---|---|---|---|
| 4 | 3 | 6131 | Cal | 89 | 100 | 1 | 3.2 | 1 | 0 | 28.8 |
| | | | Cv | NA | NA | 0.24 | 79.4 | 0.52 | -1.8 | 1.2 |
| 5 | 3 | 5469 | Cal | 91 | 100 | 1 | 3.4 | 1 | 0 | 26.8 |
| | | | Cv | NA | NA | 0.25 | 78.8 | 0.53 | -1.2 | 1.2 |
| 6 | 3 | 5249 | Cal | 91 | 100 | 1 | 3.5 | 1 | 0 | 26.8 |
| | | | Cv | NA | NA | 0.26 | 78.6 | 0.53 | -1.1 | 1.2 |
| 7 | 3 | 5168 | Cal | 91 | 100 | 1 | 3.5 | 1 | 0 | 26.7 |
| | | | Cv | NA | NA | 0.26 | 78.5 | 0.53 | -1.1 | 1.2 |
| 8 | 3 | 5143 | Cal | 91 | 100 | 1 | 3.5 | 1 | 0 | 26.7 |
| | | | Cv | NA | NA | 0.26 | 78.5 | 0.53 | -1.1 | 1.2 |
| **9** | **3** | **5140** | **Cal** | **91** | **100** | **1** | **3.5** | **1** | **0** | **26.8** |
| | | | **Cv** | **NA** | **NA** | **0.26** | **78.5** | **0.53** | **-1.1** | **1.2** |
| PPARγ | | | | | | | | | | |
| 0 | 5 | 12666 | Cal | 62 | 100 | 1.00 | 0.06 | 1.00 | 0 | 23.9 |
| | | | Cv | NA | NA | 0.38 | 1.04 | 0.34 | 0.1 | 1.3 |
| 1 | 4 | 3734 | Cal | 93 | 100 | 1.00 | 0.06 | 1.00 | 0 | 22.8 |
| | | | Cv | NA | NA | 0.47 | 0.96 | 0.39 | 0.1 | 1.4 |
| 2 | 3 | 2683 | Cal | 84 | 99 | 0.99 | 0.11 | 0.99 | 0 | 12.5 |
| | | | Cv | NA | NA | 0.65 | 0.78 | 0.49 | 0.1 | 1.7 |
| 3 | 2 | 2356 | Cal | 79 | 98 | 0.98 | 0.21 | 0.98 | 0 | 6.4 |
| | | | Cv | NA | NA | 0.71 | 0.71 | 0.54 | 0.1 | 1.9 |
| 4 | 2 | 2171 | Cal | 80 | 97 | 0.97 | 0.21 | 0.97 | 0 | 6.3 |
| | | | Cv | NA | NA | 0.76 | 0.65 | 0.57 | 0.1 | 2.1 |
| 5 | 2 | 2085 | Cal | 80 | 97 | 0.98 | 0.21 | 0.98 | 0 | 6.4 |
| | | | Cv | NA | NA | 0.77 | 0.64 | 0.57 | 0.1 | 2.1 |
| 6 | 2 | 2049 | Cal | 80 | 97 | 0.97 | 0.21 | 0.97 | 0 | 6.3 |
| | | | Cv | NA | NA | 0.77 | 0.63 | 0.57 | 0.1 | 2.2 |
| 7 | 2 | 2044 | Cal | 80 | 97 | 0.97 | 0.21 | 0.97 | 0 | 6.3 |
| | | | Cv | NA | NA | 0.77 | 0.63 | 0.57 | 0.1 | 2.2 |
| **8** | **2** | **2042** | **Cal** | **80** | **97** | **0.97** | **0.21** | **0.97** | **0** | **6.3** |
| | | | | **NA** | **NA** | **0.77** | **0.63** | **0.57** | **0.1** | **2.2** |
| ERα | | | | | | | | | | |
| 0 | 1 | 13468 | Cal | 40 | 13 | 0.13 | 0.81 | 0.13 | 0 | 1.1 |
| | | | Cv | NA | NA | -0.13 | 0.92 | -0.05 | -0.04 | 1.0 |
| 1 | 1 | 2659 | Cal | 84 | 50 | 0.50 | 0.61 | 0.50 | 0 | 1.4 |
| | | | Cv | NA | NA | -0.23 | 0.96 | 0.02 | 0.07 | 0.9 |
| 2 | 3 | 427 | Cal | 97 | 98 | 0.98 | 0.13 | 0.98 | 0 | 6.7 |
| | | | Cv | NA | NA | -0.03 | 0.88 | 0.12 | 0.09 | 1.0 |
| 3 | 1 | 179 | Cal | 82 | 81 | 0.81 | 0.38 | 0.81 | 0 | 2.3 |
| | | | Cv | NA | NA | 0.00 | 0.87 | 0.19 | 0.04 | 1.0 |
| **4** | **1** | **115** | **Cal** | **81** | **88** | **0.88** | **0.31** | **0.88** | **0** | **2.9** |
| | | | **Cv** | **NA** | **NA** | **0.24** | **0.75** | **0.22** | **0.07** | **1.2** |
| 5 | 1 | 83 | Cal | 12 | -16 | 0.91 | 0.27 | 0.91 | 0 | 3.3 |

| Model | N selected components | N features | | X cumexpvar | Y cumexpvar | R2 | RMSE | Slope | Bias | RPD |
|---|---|---|---|---|---|---|---|---|---|---|
| | | | Cv | NA | NA | 0.31 | 0.72 | 0.21 | 0.09 | 1.2 |
| 6 | Error due to multicollinearity | | | | | | | | | |
| Anti-AR | | | | | | | | | | |
| 0 | 2 | 14379 | Cal | 48 | 89 | 0.89 | 0.28 | 0.89 | 0 | 3.1 |
| | | | Cv | NA | NA | -0.20 | 0.93 | 0.12 | 0.06 | 0.9 |
| 1 | 1 | 4394 | Cal | 81 | 63 | 0.64 | 0.51 | 0.64 | 0 | 1.7 |
| | | | Cv | NA | NA | -0.26 | 0.95 | 0.11 | 0.07 | 0.9 |
| 2 | 1 | 2495 | Cal | 72 | 66 | 0.66 | 0.50 | 0.66 | 0 | 1.7 |
| | | | Cv | NA | NA | 0.02 | 0.84 | 0.18 | 0.06 | 1.0 |
| 3 | 1 | 1699 | Cal | 74 | 65 | 0.65 | 0.50 | 0.65 | 0 | 1.7 |
| | | | Cv | NA | NA | 0.30 | 0.71 | 0.29 | 0.04 | 1.2 |
| 4 | 1 | 1270 | Cal | 77 | 64 | 0.64 | 0.51 | 0.64 | 0 | 1.7 |
| | | | Cv | NA | NA | 0.45 | 0.63 | 0.37 | 0.03 | 1.4 |
| 5 | 1 | 1089 | Cal | 75 | 65 | 0.65 | 0.50 | 0.65 | 0 | 1.7 |
| | | | Cv | NA | NA | 0.51 | 0.59 | 0.40 | 0.03 | 1.5 |
| 6 | 1 | 998 | Cal | 75 | 66 | 0.66 | 0.50 | 0.66 | 0 | 1.7 |
| | | | Cv | NA | NA | 0.55 | 0.57 | 0.42 | 0.03 | 1.5 |
| 7 | 1 | 950 | Cal | 75 | 66 | 0.66 | 0.49 | 0.66 | 0 | 1.8 |
| | | | Cv | NA | NA | 0.56 | 0.56 | 0.43 | 0.03 | 1.5 |
| 8 | 1 | 926 | Cal | 75 | 67 | 0.67 | 0.49 | 0.67 | 0 | 1.8 |
| | | | Cv | NA | NA | 0.57 | 0.55 | 0.44 | 0.03 | 1.6 |
| 9 | 1 | 916 | Cal | 74 | 67 | 0.67 | 0.49 | 0.67 | 0 | 1.8 |
| | | | Cv | NA | NA | 0.58 | 0.55 | 0.44 | 0.03 | 1.6 |
| 10 | 1 | 913 | Cal | 74 | 67 | 0.67 | 0.49 | 0.67 | 0 | 1.8 |
| | | | Cv | NA | NA | 0.58 | 0.55 | 0.44 | 0.03 | 1.6 |
| **11** | **1** | **909** | **Cal** | **75** | **67** | **0.67** | **0.49** | **0.67** | **0** | **1.8** |
| | | | **Cv** | **NA** | **NA** | **0.58** | **0.55** | **0.44** | **0.03** | **1.6** |

Table S16. Receptor activity (ToxCast $AC_{50}$ values) of tentatively identified compounds detected in the optimized models of the PLS regression.

| **PLS model/activity** | **compound name** | **CAS** | **DTXSID** | **$AC_{50}$ [µM]** | **use (PubChem)** |
|---|---|---|---|---|---|
| PXR | Triphenyl phosphate | 115-86-6 | DTXSID1021952 | 0.7 | plasticizer, flame retardant |
| | Octylparaben | 1219-38-1 | DTXSID0047957 | 12.2 | |
| PPARγ | 2-Pentylfuran | 3777-69-3 | DTXSID9047679 | 4.1 | flavoring agent, fragrance |
| | Octylparaben | 1219-38-1 | DTXSID0047957 | 10 | |
| | alpha-Ionone | 127-41-3 | DTXSID0035160 | 41.5 | fragrance, odor agent |
| Anti-AR | Triphenyl phosphate | 115-86-6 | DTXSID1021952 | 9.9 | plasticizer, flame retardant |
